# A parallel cell-cycle entry pathway with inverted G1 signaling and alternate point of no return

**DOI:** 10.1101/600007

**Authors:** Chad Liu, Yumi Konagaya, Mingyu Chung, Leighton H. Daigh, Yilin Fan, Hee Won Yang, Kenta Terai, Michiyuki Matsuda, Tobias Meyer

## Abstract

Cell-cycle entry relies on an orderly progression of signaling events. To start, cells first activate the kinase cyclin D-CDK4/6, which leads to eventual inactivation of the retinoblastoma protein Rb. Hours later, cells inactivate APC/C^CDH1^ and cross the final commitment point. However, many cells with genetically deleted cyclin Ds, which activate and confer specificity to CDK4/6, can compensate and proliferate. Despite its importance in cancer, how this alternate pathway operates and whether wild-type cells use this pathway remain unknown. Here, using single-cell microscopy, we demonstrate that cells with acutely inhibited CDK4/6 enter the cell cycle with slowed and fluctuating cyclin E-CDK2 activity. Surprisingly, in this alternate pathway, the order of APC/C^CDH1^ and Rb inactivation is inverted in both cell lines and wild-type mice. Finally, we show that as a consequence of the signaling inversion, Rb inactivation replaces APC/C^CDH1^ inactivation as the point of no return. Together, we provide molecular characterization of a parallel cell-cycle entry pathway, and reveal temporal plasticity that underlies the G1 regulatory circuit.

## Introduction

To exit quiescence and start the cell cycle, non-embryonic cells first activate G1 cyclin dependent kinases (CDKs), then suppress Rb function, and finally, inactivate the E3 ubiquitin ligase APC/C^CDH1^ to trigger irreversible commitment^1,2^. In the canonical pathway, cells initiate the cell cycle by upregulating cyclin D to activate CDK4 and CDK6 (hereafter CDK4/6), resulting in eventual Rb inactivation^3,4^. In turn, Rb inactivation leads to the upregulation of critical E2F-targets such as the CDK2 activator cyclin E, APC/C^CDH1^ inhibitor EMI1, and various factors needed to replicate DNA and prevent DNA damage^5–7^. Cyclin E and EMI1 accumulation results in APC/C^CDH1^ inactivation a few hours later, which not only prepares cells metabolically for S phase, but allows for the buildup of cyclin A and DBF4 to initiate DNA replication^8–10^. Furthermore, once APC/C^CDH1^ inactivation is initiated, cell-cycle entry becomes irreversible in respect to various types of stress, reflecting an underlying G1 commitment point (Fig. 1a, top)^11–13^.

**Figure 1.**
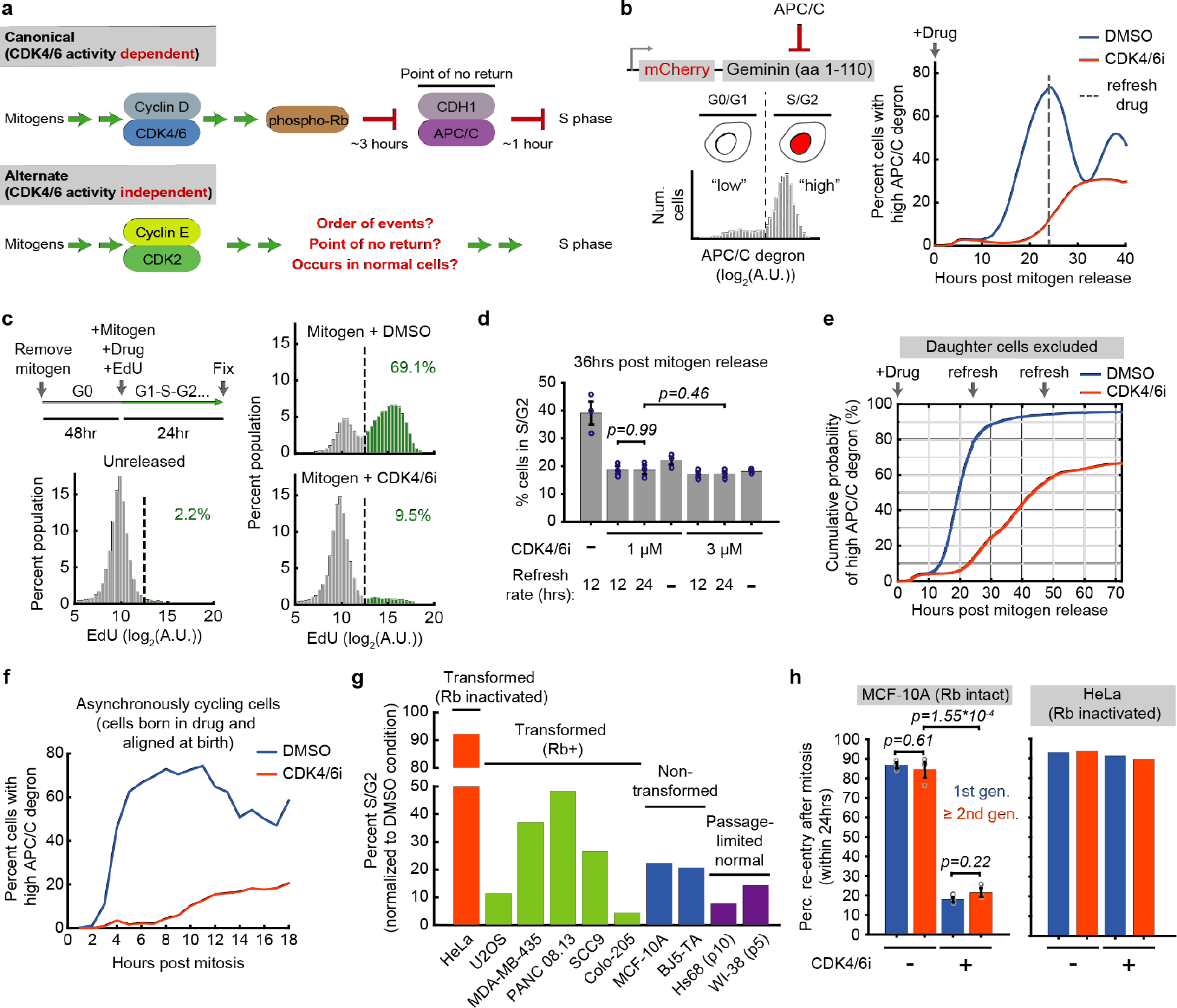
Acute CDK4/6 inhibition reveals an alternate cell-cycle entry pathway. **a**,Two paths to start the cell cycle. **b**, Left: Degron-based APC/C activity reporter. Right: Mitogen-starved MCF-10A cells expressing the APC/C activity reporter were imaged at the time of mitogen and 1μM Palbociclib (CDK4/6 inhibitor) addition. Time points taken every 12 minutes. CDK4/6 inhibitor refreshed at 24hrs. 1 of n=5 biological replicates. **c**, MCF-10A cells mitogen-released in the presence of 100nM EdU with or without CDK4/6 inhibitor. Cells were fixed after 24hrs (26838, 36549, 29371 cells for unreleased, DMSO, and CDK4/6 inhibitor; 1 of n=3 biological replicates). **d**, Cells were mitogen-released with or without indicated CDK4/6 inhibitor concentrations and the drug was refreshed every 12hrs, 24hrs, or not at all. Cells fixed at 36hrs after mitogen release. S/G2 cells determined via EdU incorporation and DNA content. Error bars: SEM from 3 biological replicates; p-values calculated using two-sided, two-sample t-tests. **e**, CDF of mitogen-released cells inactivating APC/C^CDH1^. Only cells present at the time of mitogen-release are tracked. **f**, APC/C activity in cells born into DMSO or CDK4/6 inhibitor. Cells aligned at birth. 1 of n=3 biological replicates. **g**, Different cell lines treated with 1μM CDK4/6 inhibitor for 48hrs and refreshed at 24hrs. Percent S/G2 defined as >2n DNA (determined by Hoechst) and high endogenous geminin (an APC/C^CDH1^ substrate). Percentages normalized to the DMSO-treated condition. **h**, Generation comparison of cell-cycle re-entry percentages in MCF-10A (Rb intact) and HeLa (Rb inactivated) cells (error bars: SEM from 3 biological replicates; p-values calculated using two-sided, two-sample t-tests).

When D-class cyclins are genetically ablated, however, many non-transformed cells can still start the cell cycle by directly activating cyclin E-CDK2^14^. Notably, deletion of both D and E-class cyclins blocks cell-cycle progression in MEFs and most cell lineages in a developing embryo^15^. These results suggest that in many non-transformed cells, E-class cyclins can compensate if D-class cyclins are missing. While the canonical cyclin D-CDK4/6 initiated pathway has been extensively characterized in a variety of contexts, the alternate cyclin E-CDK2 initiated pathway has mainly been characterized using knockout models and in cancer, where cells can bypass CDK4/6 inhibition via c-Myc upregulation of cyclin E-CDK2 activity, amplification of cyclin E, or downregulation of CDK2 inhibitors^16–23^. However, since the cyclin E-CDK2 initiated pathway has only been described in cancer and in scenarios where cyclin Ds are deleted at germline, it remains unknown whether wild-type cells can make use of this alternate pathway. Furthermore, it is generally assumed that cell-cycle entry in this alternate pathway requires a set order of events where cyclin E-CDK2 activation is followed by Rb inactivation, which is then followed by APC/C^CDH1^ inactivation and irreversible commitment. This hypothesis of a rigid order underlying G1 progression has also not been experimentally tested (Fig. 1a, bottom).

Here, applying live and fixed single-cell microscopy, we show that cells with acutely inhibited CDK4/6 activity still proliferate but with cyclin E-CDK2 being initially activated non-persistently and without Rb hyperphosphorylation. Surprisingly, the order of Rb and APC/C^CDH1^ inactivation is inverted both in non-transformed cell lines as well as the small intestinal crypts of wild-type mice. Thus, our study argues that the alternate pathway can be employed under normal conditions and also argues against a rigid order of signaling events in G1. Finally, we show that this signaling inversion leads to a point of no return that is marked by Rb inactivation instead of APC/C^CDH1^ inactivation. Thus, to start the cell cycle, cells first activate CDKs via upregulation of D or E-class cyclins, then inactivate Rb and APC/C^CDH1^ in an interchangeable manner, and finally, commit to the cell cycle only after both Rb and APC/C^CDH1^ are inactivated.

## Results

### Acute CDK4/6 inhibition reveals an alternate cell-cycle entry pathway

To monitor cell-cycle entry at the single-cell level, we stably transduced non-transformed MCF-10A epithelial cells with a previously characterized APC/C^CDH1^ activity reporter that is degraded during G0/G1 and synthesized during late G1/S/G2 phase (Fig. 1b, left)^12,24^. We deprived cells of growth factors for 48 hours, and then added back mitogen in the presence or absence of the specific CDK4/6 inhibitor Palbociclib^25,26^ (hereafter just CDK4/6 inhibitor) along with EdU, a nucleoside analog that is incorporated by cells in S phase. Markedly, we found that a fraction of CDK4/6 inhibitor treated cells still inactivated APC/C^CDH1^ and incorporated EdU (Fig. 1b, right; Fig. 1c), arguing that cells can enter S phase without CDK4/6 activity even when cyclin Ds are not deleted at germline. Entry into the cell cycle was independent of the drug refreshing rate and was also not further inhibited by a three-fold increase in inhibitor concentration (Fig. 1d). Cell-cycle entry in CDK4/6-inhibited cells was notably delayed and reduced compared to cells with intact CDK4/6 activity, demonstrating an extended G1 phase. However, when tracking cells for three days, we found that a majority of them eventually inactivated APC/C^CDH1^ (Fig. 1e). Since MEFs genetically depleted of cyclin Ds also exhibit less efficient entry into S phase as well as a longer G1^14^, we reasoned that acute chemical inhibition of CDK4/6 phenocopies the ablation of cyclin Ds, suggesting that the alternate pathway is not only a long-term compensation mechanism induced by germline cyclin D loss.

We next broadened the analysis to asynchronously cycling cells, where most newborn daughter cells do not enter quiescence and start preparing and committing to the cell cycle in mother cells^1,27–29^. By *in silico* aligning cells at the time of birth, we found the same extended G1 phase also in CDK4/6-inhibited cycling cells (Fig. 1f, Extended Data Fig. 1a). Furthermore, acute CDK4/6 inhibitor treatment reduced the percentage of cells in S/G2 in a variety of normal and cancer cells, consistent with CDK4/6 inhibition mediating an extended G1 phase (Fig. 1g). We note that cells need intact Rb to be responsive to CDK4/6 inhibition, explaining the lack of effect of CDK4/6 inhibition in HeLa cells (Fig. 1g)^22^. Finally, even though sister cells mostly had the same fate after birth (Extended Data Fig. 1b), live-cell analysis showed that subsequent generations of CDK4/6-inhibited cells were not significantly more likely to re-enter the cell cycle compared to the first generation (experiment setup: Extended Data Fig. 1c; results: Fig. 1h). This lack of correlation across generations suggests that it is not a clonal subpopulation of MCF-10A cells that gained heritable resistance to CDK4/6 inhibition.

### Cells without CDK4/6 activity exhibit slowed and fluctuating cyclin E-CDK2 activity

Because cells without CDK4/6 activity rely on cyclin E-CDK2 for cell-cycle entry, we stably expressed in the same cells a previously characterized live-cell cyclin E/A-CDK activity reporter based on a DHB peptide (Fig. 2a)^30–32^. This reporter measures relative kinase activity as the ratio of the cytoplasmic over nuclear reporter concentrations. The activity ratio increases from approximately 0.5 at basal, to 1 at the G1/S transition, and to 1.5 when cells enter mitosis. We note that the reporter detects primarily cyclin E-CDK2 activity in G1 since there is minimal cyclin A protein or CDK1 activity in G1^12,31,33,34^. Starting live-cell imaging at the time of mitogen release to determine an activity baseline, we found that a portion of CDK4/6 inhibitor-treated cells indeed upregulated cyclin E/A-CDK activity (Fig. 2b, right panel, blue cells, defined as >0.7 on the last time point). Activation in these cells was delayed (Extended Data Fig. 2a) and surprisingly, the kinetics were distinct from that of control cells, where cyclin E/A-CDK activity in most cells increased in a persistent and linear fashion (Fig. 2b, left). CDK4/6-inhibited cells built up cyclin E/A-CDK activity slower (Fig. 2b, blue cells) and had fluctuating activities (green cells, defined as <0.7 at 24hr, but >0.7 at least once between 8 and 24hrs). Control experiments with a DHB mutant that cannot translocate showed that most fluctuations were not measurement noise, and that the fluctuating cells had not started S phase (Extended Data Fig. 2b-c). The increase in cyclin E/A-CDK activity was regulated by cyclin E and the CDK2 inhibitor p21 (Fig. 2c), suggesting that the CDK activity fluctuations and increase result from competition between dynamically changing cyclin E or p21 levels. Because stress is known to upregulate p21 and downregulate CDK2 activity^12,35^, we hypothesize that cells in the alternate pathway are more sensitive to stress given its slow and fluctuating CDK2 activity. Indeed, a fifteen-minute pulse of neocarzinostatin (can cause DNA breaks in quiescence^36^) prior to mitogen release confirmed that CDK2 activity in cells without CDK4/6 activity is proportionally more affected (Fig. 2d).

**Figure 2.**
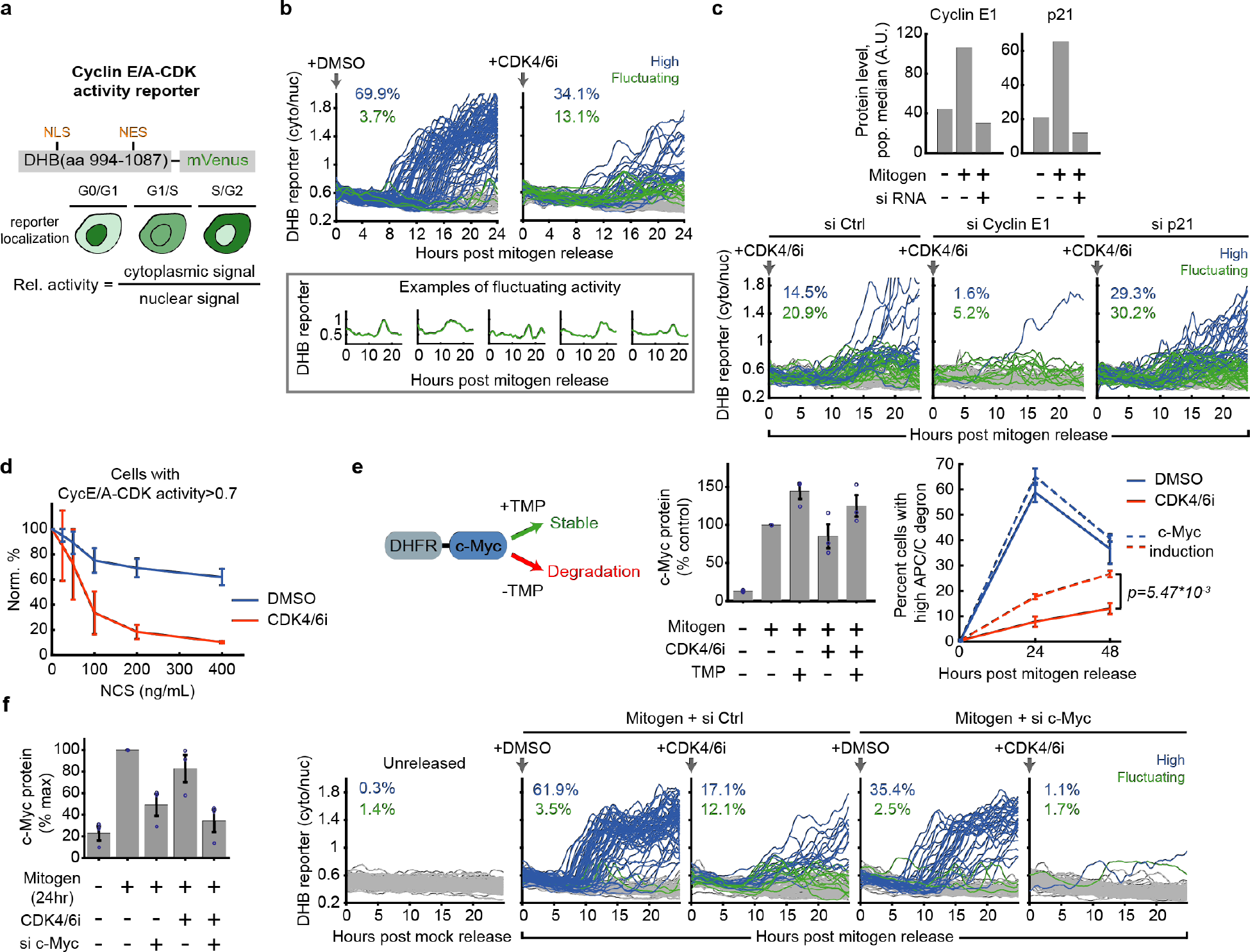
Cells without CDK4/6 activity exhibit fluctuating cyclin E-CDK2 activity. **a**, DHB-based cyclin E/A-CDK activity reporter. **b**, Single-cell traces of cyclin E/A-CDK activity after mitogen release with or without 1μM CDK4/6 inhibitor (100 cells plotted; 2782 and 8538 cells total for DMSO and CDK4/6 inhibitor; 1 of n=2 biological replicates). Activity baseline at time 0 is determined to be at most 0.7. Blue: >0.7 at last time point. Green: <0.7 at last time point, exceeded 0.7 at some point between 8 and 24hrs. Bottom: Examples of fluctuating cyclin E/A-CDK activity. **c**, Top: immunofluorescence validation of cyclin E1 and p21 knockdown. Bottom: effect of knockdown on cyclin E/A-CDK activity in cells treated with CDK4/6 inhibitor. Traces classified the same way as **b**(100 cells plotted; 2958, 3707, 3113 cells for si Ctrl, si Cyclin E1, and si p21; 1 of n=2 biological replicates). **d**, Cells were treated with neocarzinostatin (NCS) for 15min, and then mitogen-released for 24hrs. Percent cells with high cyclin E/A-CDK activity (defined as reporter ratio>0.7) are displayed. Percentages normalized to 0ng/mL condition. Error bars: standard deviation of 2 biological replicates. **e**, Cells were stably infected with DHFR-Myc that is unstable until TMP molecule addition. c-Myc levels were induced to ~50% above control after adding 10μM TMP at the time of mitogen release. Cells fixed at 24 and 48hrs after mitogen release. Drugs refreshed every 24hrs. Error bars: SEM from 3 biological replicates; p-values calculated using two-sided, two-sample t-tests. **f**, Left: immunofluorescence of c-Myc knockdown validation (error bars: SEM from 3 biological replicates). Right: cyclin E/A-CDK activity after mitogen-release in MCF-10A (100 cells plotted, 569, 3446, 3904, 3935, 3767 total cells for unreleased, si Ctrl+DMSO, si Ctrl+CDK4/6 inhibitor, si c-Myc+DMSO, and si c-Myc+CDK4/6 inhibitor, respectively), representative of n=5 biological replicates.

Similar to previous reports that c-Myc overexpression can promote S-phase entry in cells where CDK4/6 activity was inhibited by p16 expression^16^, a 50% increase of c-Myc levels was sufficient to significantly increase the fraction of CDK4/6-inhibited cells entering the cell cycle (Fig. 2e). Furthermore, c-Myc knockdown in the presence of CDK4/6 inhibitor reduced cyclin E transcription and abolished cyclin E-CDK2 activity and S-phase entry (Extended Data Fig. 3a, Fig. 2f, Extended Data Fig. 3b). We conclude that c-Myc and cyclin D-CDK4/6 activities are driving forces that can promote cell-cycle entry through two different paths - one with slowed and gradual CDK2 activation, and one with fast and persistent CDK2 activation.

### Cyclin E-CDK2 is activated without Rb hyperphosphorylation in cells lacking CDK4/6 activity

To understand the role of Rb inactivation in cells with inhibited CDK4/6 activity, we examined the phosphorylation status of Rb, a key repressor of E2Fs. When phosphorylated on most or all of its accessible sites (“hyperphosphorylation”), Rb releases E2F transcription factors to upregulate cyclin E (Fig. 3a)^4–6,37^. To measure Rb phosphorylation, we first restricted our live-cell analysis to G1 cells at early stages of CDK2 activation by selecting cells that had not yet inactivated APC/C^CDH1^ and had just activated cyclin E-CDK2. We then fixed cells and measured the phosphorylation status of serine 807 and serine 811 (S807/S811) on Rb by normalizing the immunofluorescent phosphorylation signal against that of total Rb in each cell (Fig. 3b). This analysis revealed that DMSO-treated control cells that had just activated cyclin E-CDK2 already had S807/S811 phosphorylated at levels nearly identical to that of S/G2 cells, when Rb is hyperphosphorylated (Fig. 3c, top and middle)^37^. However, unexpectedly, in CDK4/6 inhibited cells, we observed a much lower signal of phospho-S807/S811 even though the measured cyclin E/A-CDK activity level was the same (Fig. 3c, bottom). Control experiments showed the same result of suppressed Rb phosphorylation in CDK4/6 inhibitor treated BJ-5ta, a non-transformed human foreskin fibroblast (Fig. 3d), as well as when we used two alternative CDK4/6 inhibitors with different pharmacological properties (Extended Data Fig. 4). Since Rb hyperphosphorylation and inactivation requires most or all sites to be phosphorylated^37^, these results indicate that early cyclin E-CDK2 activation in CDK4/6-inhibited cells does not require Rb hyperphosphorylation and is therefore the result of direct upregulation of cyclin E expression by c-Myc. Furthermore, this data suggests that a key role of CDK4/6 is to confer persistence to cyclin E-CDK2 activation in G1. We note that since Rb is the best characterized substrate of CDK4/6^3,4,37^, the suppressed Rb phosphorylation also demonstrates that cyclin D-CDK4/6 complexes were inhibited in the cells undertaking the alternate pathway.

**Figure 3.**
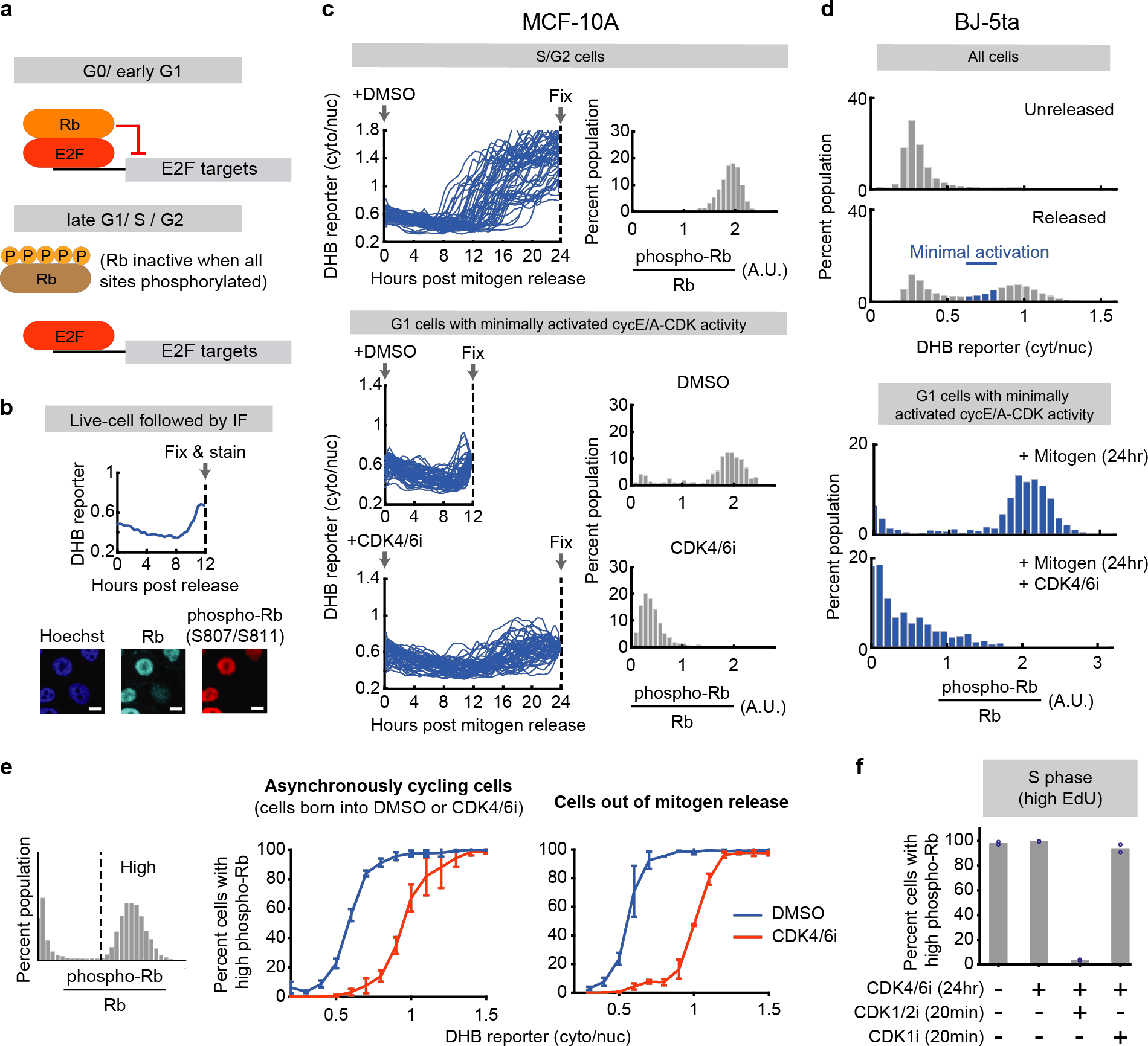
Cyclin E-CDK2 is activated without Rb hyperphosphorylation in cells lacking CDK4/6 activity. **a**, E2F is active when Rb is hyperphosphorylated. **b**, Phospho-Rb analysis after live-cell tracking of cyclin E/A-CDK activity. Scale bar=10μm. **c**, Phospho-Rb(S807/S811) analysis of cells that have inactivated APC/C^CDH1^ (top; 50 cells plotted, 2315 cells total), and cells that have initiated cyclin E/A-CDK activity and have not inactivated APC/C^CDH1^ (bottom; 50 cells plotted, 364, 1023 cells for DMSO and CDK4/6 inhibitor). 1 of n=3 biological replicates. **d**, BJ-5ta cells expressing the cyclin E/A-CDK activity reporter and APC/C degron reporter were mitogen-released with or without 1μM CDK4/6 inhibitor, fixed after 24hrs, and stained for Rb and phospho-Rb(S807/S811). G1 cells with recently activated cyclin E/A-CDK activity were analyzed. **e**, Cells were grouped into high and low phospho-Rb(S807/S811) signal populations. The percentages of cells with high signal were then determined for each bin of the cyclin E/A-CDK activity. Middle: Cells born into DMSO or CDK4/6 inhibitor. Error bars denote standard deviation of 3 biological replicates. Right: Cells mitogen-released with DMSO or CDK4/6 inhibitor. Error bars denote standard deviation of 2 biological replicates. **f**, Asynchronously cycling cells treated with or without CDK4/6 inhibitor for 24hrs. Cells were then incubated with 10μM EdU for 15min, and then treated with DMSO, CDK1/2i (3μM), or CDK1i (10μM) for 20min. Cells were then fixed and high EdU signal cells were examined. Data points denote biological replicates.

Markedly, when we plotted the percentages of cells with high phospho-Rb (S807/S811) signal as a function of cyclin E/A-CDK activity, we found that CDK4/6-inhibited cells did eventually reach the high phospho-Rb state, but only at much higher cyclin E/A-CDK activity levels (Fig. 3e). These results suggest that Rb inactivation may occur much later in the alternate pathway. This delay is likely due to CDK4/6’s ability to contribute to Rb phosphorylation in the G1 canonical pathway^3,4,37^, and also argues for the first time that much higher cyclin E/A-CDK activity is required in CDK4/6-inhibited cells to reach maximal Rb phosphorylation. When we gated for CDK4/6 activity-independent cells already in S phase and measured phospho-Rb signal after treatment with the CDK1/2 and CDK1 inhibitors, the high phospho-Rb signal disappeared after incubation with CDK1/2 inhibitor but not CDK1 inhibitor (Fig. 3f). Together, we conclude that in CDK4/6-inhibited cells, Rb is phosphorylated only in late G1 at much higher levels of cyclin E/A-mediated CDK2 activity compared to cells that activate cyclin D-CDK4/6.

### The canonical signaling order of Rb inactivation before APC/C^CDH1^ inactivation is inverted in the alternate pathway

Due to the need for high CDK2 activity for Rb phosphorylation in the alternate pathway (Fig. 3e), we hypothesized that, in the presence of CDK4/6 inhibitor, full Rb phosphorylation and inactivation may occur only after APC/C^CDH1^ inactivation. This is possible since APC/C^CDH1^ inactivation can be mediated by cyclin E-CDK2 phosphorylation on CDH1 even in the absence of EMI1^12,13^. This hypothesis is further strengthened by the finding that APC/C^CDH1^ inactivates at approximately the same level of cyclin E/A-CDK activity in the alternate and canonical pathway (Extended Data Fig. 5a). To directly determine the order of events, we measured high phospho-Rb(S807/S811) signal in cells that had just inactivated APC/C^CDH1^ using the increase in the APC/C degron signal as a proxy for time after APC/C^CDH1^ inactivation (Fig. 4a, bottom left histograms). Such a conversion from concentration to time is possible since the concentration of the APC/C degron reporter increases in a linear fashion after APC/C^CDH1^ inactivation. Consistent with our previous results, most of the DMSO-treated cells that just inactivated APC/C^CDH1^ already had high phospho-Rb(S807/S811) signal (Fig. 4a, DMSO histograms). Strikingly, in CDK4/6-inhibited cells, the high Rb phosphorylation signal peak in the histogram only started to appear after a delay following the buildup of APC/C degron (Fig. 4a, CDK4/6i histograms). The same delay also occurs in asynchronously cycling cells (Extended Data Fig. 5b). This argues that the order of Rb inactivation and APC/C^CDH1^ inactivation is inverted in CDK4/6-inhibited cells with Rb inactivation occurring after APC/C^CDH1^ inactivation.

**Figure 4.**
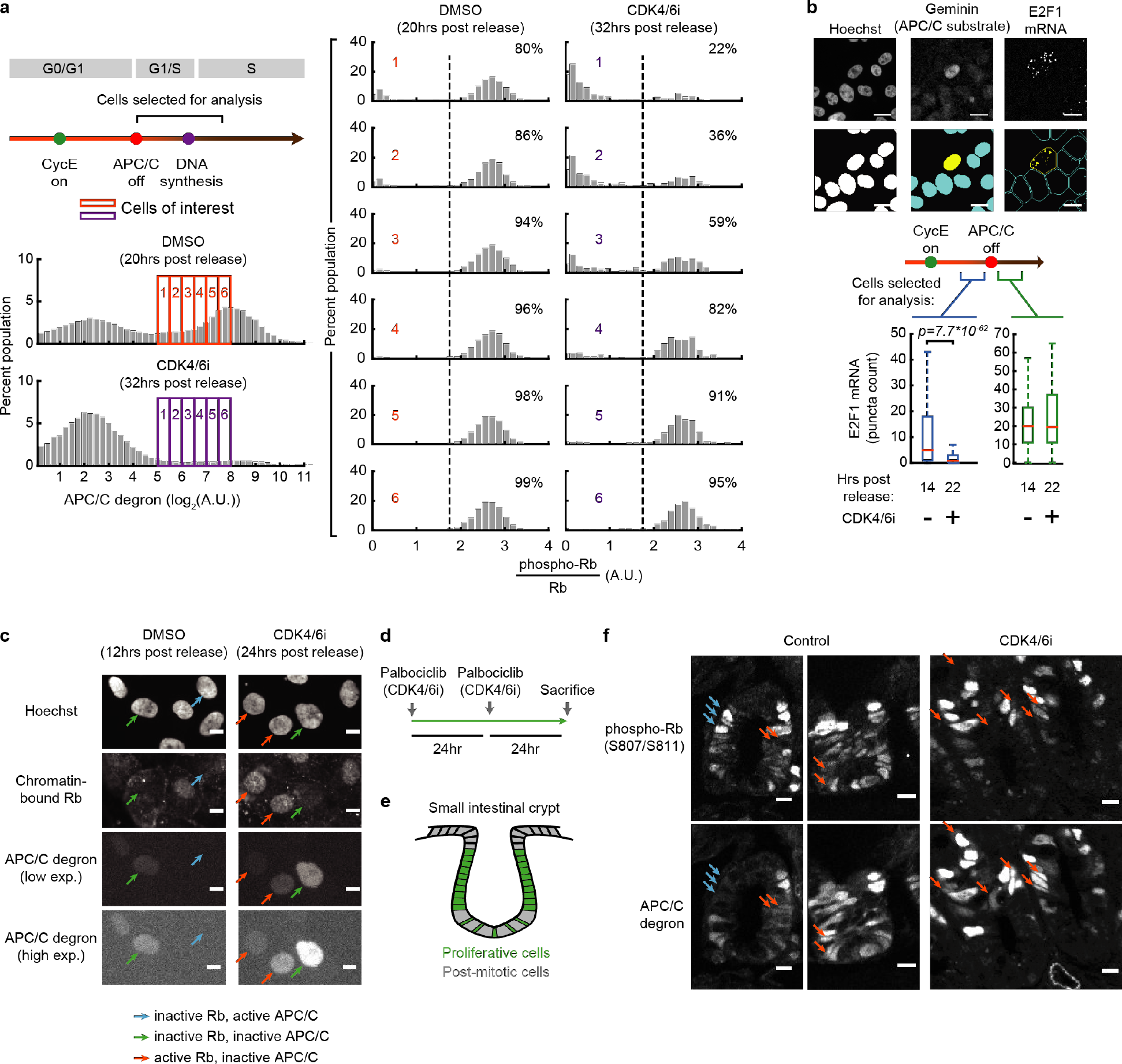
The canonical signaling order of Rb inactivation before APC/C^CDH1^ inactivation is inverted in the alternate pathway. **a**, Cells mitogen-released with DMSO or 1μM CDK4/6 inhibitor and have recently inactivated APC/C^CDH1^ were analyzed for phospho-Rb(S807/S811) signal (>800 cells per histogram for DMSO, >200 cells per histogram for CDK4/6 inhibitor; 1 of n=3 biological replicates). **b**, Top: Single-cell mRNA quantification (scale bar=10μm). Bottom: E2F1 mRNA in cells released with or without CDK4/6 inhibitor. High cyclin E/A-CDK activity defined as ratio of signal>0.7. p-values calculated using two-sided, two-sample t-tests (from left to right: n=784, 1484, 638, and 98 cells). 1 of n=3 biological replicates. **c**, Mitogen-released cells were assayed for chromatin-bound Rb. Green arrows: cells that have inactivated both Rb and APC/C^CDH1^; blue arrows: cells that have inactivated Rb but not APC/C^CDH1^; orange arrows: cells that have inactivated APC/C^CDH1^ but not Rb. Scale bar: 10μm. 1 of n=2 biological replicates. **d**, Experiment setup for drugging mice. **e**, Illustration of proliferative and post-mitotic cells in the small intestinal crypt. **f**, Mice small intestinal crypts that were treated with vehicle or CDK4/6 inhibitor. The cells express the APC/C activity reporter and were stained for phospho-Rb (S807/S811). Blue arrows denote example cells with high phospho-Rb and low APC/C degron signal. Orange arrows denote cells with high APC/C degron and low phospho-Rb signal. Crypts are oriented in an U-shape.

To directly measure whether E2F targets are also only activated after APC/C^CDH1^ inactivation in the alternate pathway, we measured the transcripts of E2F1, a well-characterized E2F target that is suppressed by Rb^6^. Using single-cell mRNA FISH analysis, we confirmed that E2F1 transcription was suppressed by CDK4/6 inhibition in cells with high cyclin E-CDK2 activity and undetectable APC/C degron levels (Fig. 4b, left; Extended Data Fig. 5c, left). However, only after a delay following APC/C^CDH1^ inactivation did we observe a significant increase in E2F1 transcription in CDK4/6-inhibited cells (Fig. 4b, right; Extended Data Fig. 5c, right).

To further validate that Rb is indeed inactivated only after APC/C^CDH1^ inactivation, we made use of previous findings that inactivated Rb dissociates from the chromatin^38^. We thus measured pre-extracted/chromatin-bound Rb in single cells^39^. Control experiments confirmed that in the canonical pathway, Rb dissociates from the chromatin (low chromatin-bound Rb signal) prior to APC/C^CDH1^ inactivation and remains dissociated after APC/C^CDH1^ is turned off (Fig. 4c, DMSO condition, green and blue arrows; Extended Data Fig. 6a). However, in CDK4/6-inhibited cells, Rb only dissociated from the chromatin after a delay and after significant APC/C degron buildup (Fig. 4c, CDK4/6 inhibitor condition, orange and green arrows; Extended Data 6b-c). Together, our results of delayed Rb phosphorylation, Rb chromatin-dissociation, and E2F1 activation demonstrate that CDK4/6-inhibited cells have an alternate, inverted G1 pathway, with APC/C^CDH1^ being inactivated prior to Rb inactivation.

While our experiments thus far identified a markedly different cell-cycle entry sequence, we have not determined if such order operates in normal physiological settings. We thus treated adult mice expressing the APC/C^CDH1^ activity reporter^40^ with CDK4/6 inhibitor for 48 hours and examined the small intestinal crypts, a proliferative region where cells are continuously replenished (Fig. 4d-e)^41^. To distinguish cells in the alternate or canonical pathway, we stained the crypts for phospho-Rb (S807/S811) and compared the signal to that of the APC/C reporter. We note that this approach can only distinguish between the two pathways in the subset of cells in mid-G1 phase where the alternate pathway is marked by high APC/C degron and low phospho-Rb signals, and the canonical pathway by low APC/C degron and high phospho-Rb signals (Fig. 4f, control condition, examples of canonical cells are denoted by blue arrows). In this analysis, most cells in the population are in early G1 or G0, marked by low APC/C degron expression and low Rb phosphorylation, or in late G1, S or G2, marked by increased APC/C degron expression and phosphorylated Rb. Markedly, this analysis showed that even in normal conditions, crypt cells can be observed to have either the canonical or alternate sequence (Fig. 4f, control condition, orange arrows). To quantify these observations, we used post-mitotic cells as negative controls to unbiasedly select thresholds for phosho-Rb and APC/C degron positive cells, and calculated the percentage of cells with high phospho-Rb signal, low APC/C degron signal or vice versa (Extended Data Fig. 7a, blue and orange). The quantification further confirms that both alternate and canonical paths can be found in crypt cells under normal unperturbed conditions. Furthermore, control experiments showed that systemic CDK4/6 inhibition lowers the percentage of cells in the canonical pathway, as predicted, but does not decrease the percentage of cells undertaking the alternate sequence (Extended Data Fig. 7b, CDK4/6 inhibitor condition; Fig. 4f). Thus, normal cells have two options and can enter the cell cycle by either taking the canonical cyclin D-CDK4/6 initiated path, or the alternate cyclin E-CDK2 initiated path.

### APC/C^CDH1^ inactivation is reversible until Rb inactivation in cells lacking CDK4/6 activity

The inversion of Rb and APC/C^CDH1^ inactivation in CDK4/6-inhibited cells suggest that Rb inactivation might be a new rate-limiting step in G1 progression with cells now waiting for Rb instead of APC/C^CDH1^ inactivation to commit to the cell cycle and enter S phase. If true, the point of no return, or commitment point, for cell-cycle entry would be a coincidence detection mechanism representing an interchangeable order of APC/C^CDH1^ and RB inactivation rather that a predetermined order of events. Indeed, by plotting phospho-Rb(S807/S811) signal against incorporated EdU signal, we confirmed that Rb is phosphorylated prior to DNA incorporation (Fig. 5a). We found that in both mitogen released and asynchronously cycling cells, Rb is phosphorylated ≥4 hours after APC/C^CDH1^ inactivation, and DNA is synthesized ≥6 hours after APC/C^CDH1^ inactivation as opposed to 1 hour in the canonical pathway (Fig. 5b and Extended Data Fig. 8a). The long delay from APC/C^CDH1^ inactivation to S phase suggests that cells without CDK4/6 activity wait for Rb inactivation and E2F activation instead of APC/C^CDH1^ inactivation to start replicating their DNA.

Previous studies of the canonical pathway showed that APC/C^CDH1^ inactivation can be initiated either by E2F-mediated activation of cyclin E-CDK2 or expression of the APC/C^CDH1^ inhibitor EMI1^12,13^. However, irreversible APC/C^CDH1^ inactivation requires a double negative feedback between EMI1 and APC/C^CDH1^ (Fig. 5c)^13^. In cell-cycle control, irreversible steps are necessary for maintaining the proper order of cell-cycle phases^42–44^, and without EMI1-mediated APC/C^CDH1^ inactivation, genome integrity is compromised^11,45,46^. As cell can temporally inactivate cyclin E/A-CDK in response to DNA damage, a role of EMI1 is likely to prevent inadvertent APC/C^CDH1^ reactivation in S phase, where endogenous DNA damage can suppress cyclin E/A-CDK activity^34^. In CDK4/6-inhibited cells, E2F-mediated EMI1 transcription is delayed similar to that of E2F1 (Extended Data Fig. 8b). Consistent with a lack of EMI1, which results in slower APC/C^CDH1^ inactivation kinetics^12,13^, we found slowed APC/C^CDH1^ inactivation in CDK4/6-inhibited cells (Extended Data Fig. 8c-d). We next directly tested for reversibility of APC/C^CDH1^ inactivation. Unlike cells with active CDK4/6 (Fig. 5d), cells without CDK4/6 activity reactivated APC/C^CDH1^ after CDK1/2 inhibition (Fig. 5e, gray box). APC/C^CDH1^ inactivation only becomes irreversible several hours later (Fig. 5e, right panels), and knock down of EMI1 at this time point reactivates APC/C^CDH1^ (Extended Data Fig. 8e-f). Thus, APC/C^CDH1^ inactivation is initially reversible in CDK4/6-inhibited cells until Rb is inactivated and EMI1 can be synthesized.

**Figure 5.**
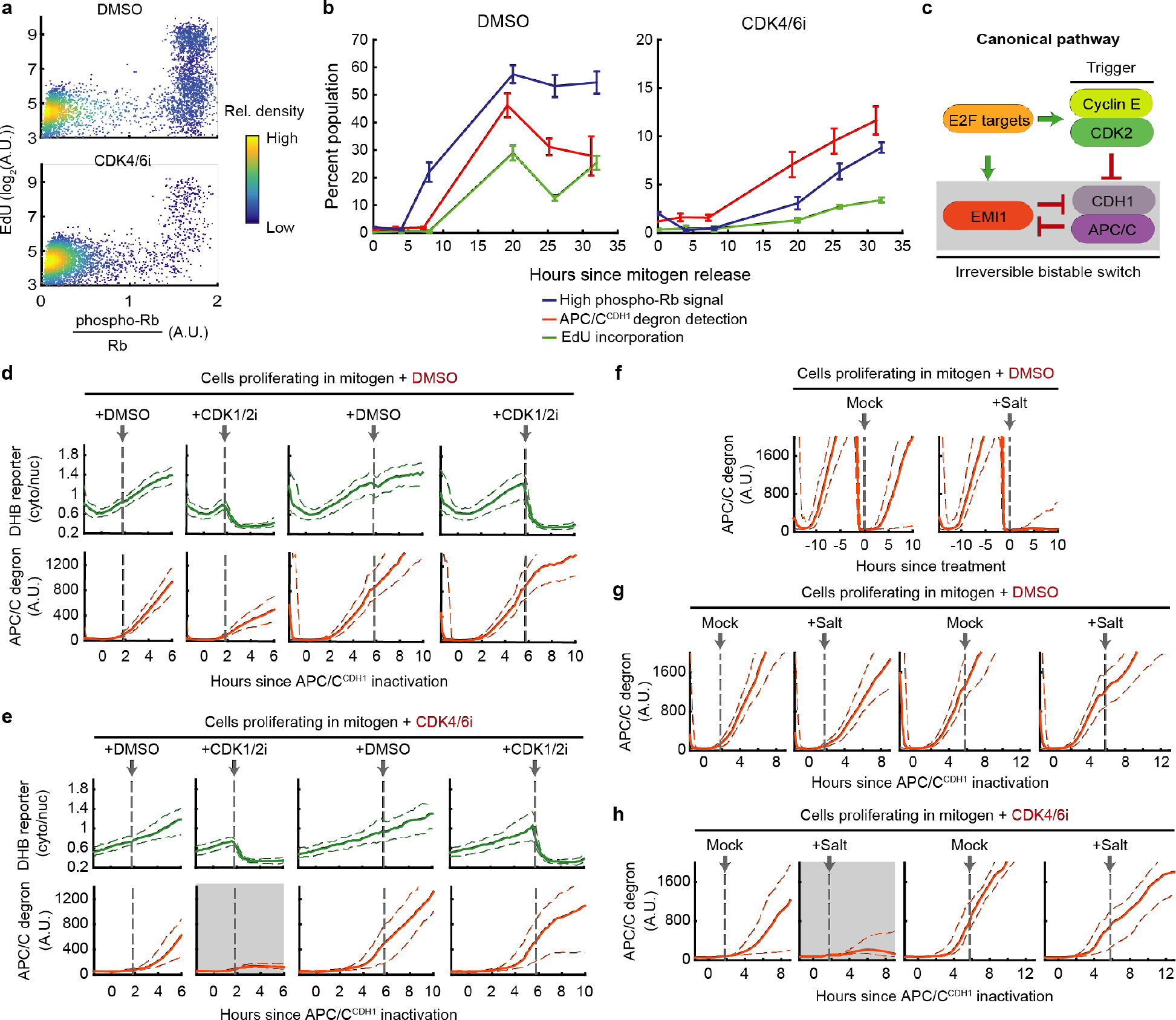
APC/C^CDH1^ inactivation is reversible until Rb inactivation in cells lacking CDK4/6 activity. **a**, Cells born into DMSO or 1μM CDK4/6 inhibitor were treated with 10μM EdU for 15min, and then assayed for EdU and phospho-Rb(S807/S811) (5000 cells plotted, 1 of n=3 biological replicates). **b**, Mitogen-released cells treated with DMSO or CDK4/6 inhibitor were measured for percent cells with high phospho-Rb(S807/S811) signal, high APC/C degron, and high EdU signal (after 15min of 10μM EdU treatment). Error bars: SEM from 3 biological replicates. **c**, EMI1 ensures irreversible APC/C^CDH1^ inactivation. **d-h**, Cells born into DMSO or CDK4/6 inhibitor and have inactivated APC/C^CDH1^ (with the exception of **f**, where it is just cells born into DMSO or CDK4/6 inhibitor) were treated with 3μM CDK1/2i (**d-e**) or an extra 100mM salt (**f-h**). Dashed lines denote 25^th^ and 75^th^ percentile. Gray boxes denote conditions where cells re-activated APC/C^CDH1^. n=2 biological replicates; >30 cells per condition. In **g-h**, to enrich for Rb inactivated cells in the 6hrs post APC/C^CDH1^ inactivation conditions, only cells with cyclin E/A-CDK>0.8 right before treatment were analyzed (threshold determined from Figure 3e).

APC/C^CDH1^ inactivation is an important step in canonical cell-cycle entry as cells remain sensitive to stress in the canonical pathway until but not after APC/C^CDH1^ inactivation (for hypertonic stress, Fig. 5f-g; for oxidative stress, Extended Data Fig. 8g-h), arguing that this step represents the commitment point^12^. We thus tested for commitment in respect to stress in the alternate pathway. Markedly, in asynchronously cycling CDK4/6-inhibited cells, stress applied two hours after APC/C^CDH1^ inactivation resulted in reactivation of APC/C^CDH1^ (Fig. 5h and Extended Data Fig. 8i, gray boxes). We next determined if Rb inactivation coincides with a loss of stress sensitivity in the alternate pathway by examining cells that were (1) treated with stress six hours after APC/C^CDH1^ inactivation, and (2) had cyclin E/A-CDK activity>0.8 at the time of treatment. The additional CDK activity gating (threshold determined from Figure 3e) enriches for cells with inactivated Rb at the time of stress treatment. Applying these gating conditions, we found that APC/C^CDH1^ inactivation indeed became irreversible in respect to stress (Fig. 5h; Extended Data Fig. 8i, right panels). Together with the EMI1 results, we conclude that in cells proliferating without CDK4/6 activity, the commitment point that is canonically associated with APC/C^CDH1^ inactivation is now delayed and marked instead by Rb inactivation.

Given this unexpected inversion of signaling order and commitment process, we tested for potential cell-cycle defects in the alternate pathway. We found that these cells exhibited similarly low S/G2 phase DNA damage compared to the canonical pathway as measured by γH2A.X and 53BP1 signals (Extended Data Fig. 9a-b). By measuring the amount of time between APC/C^CDH1^ inactivation and mitosis, we also did not find prolonged S and G2 phases when considering the five-hour delay between APC/C^CDH1^ inactivation and DNA synthesis described in Extended Data Figure 8a (Extended Data Fig. 9c). Based on these results, it is unlikely that the S and G2 phases of the cell cycle are severely affected by inversion of Rb and APC/C^CDH1^ inactivation or the change in point of no return. Combined with the small intestinal crypt data, our data suggests that the alternate sequence is a viable path for cell-cycle entry.

## Discussion

Seminal genetic work established that cells can proliferate without D-class cyclins, which normally activate and confer specificity to CDK4/6^14^. In CDK4/6 knockout cells, cyclin Ds may still contribute to cell-cycle entry regulation by activating other CDKs^47^. In the case of ablated cyclin Ds, the adaptability is due to activation of cyclin E-CDK2, though this alternative pathway of cyclin E-CDK2 activation without cyclin D-CDK4/6 activation has not yet been demonstrated to operate in wild-type cells (we note that genetic mutations may not reflect acute depletions^48,49^). Using single cell microscopy techniques, our study demonstrates that wild-type cells can use cyclin E-CDK2 instead of cyclin D-CDK4/6 to start the cell cycle, though in a non-persistent and delayed fashion. Furthermore, in the alternate cyclin E-CDK2-initiated pathway, we found that cells invert the order of Rb and APC/C^CDH1^ inactivation, and only cross the point of no return after Rb is inactivated instead of APC/C^CDH1^ inactivation in the canonical pathway (Figure 6). Our study not only provides detailed molecular characterization of this parallel pathway, it demonstrates that the pathway can operate in normal cells in vitro and in vivo – providing evidence for temporal signaling plasticity in G1 without a rigid order of events. Given that the point of no return for the cell cycle is only triggered in the alternate pathway after Rb has also been inactivated, our study argues that cell-cycle commitment requires that both Rb and APC/C^CDH1^ to be inactivated independent of the sequence of events. The characterization of such an alternate cyclin E-CDK2-initiated pathway is broadly relevant as it may have main or backup roles in regulating proliferation in a variety of biological contexts, and the pathway may also have a role in controlling tissue replacement rates due to its much longer G1 phase.

**Figure 6.**
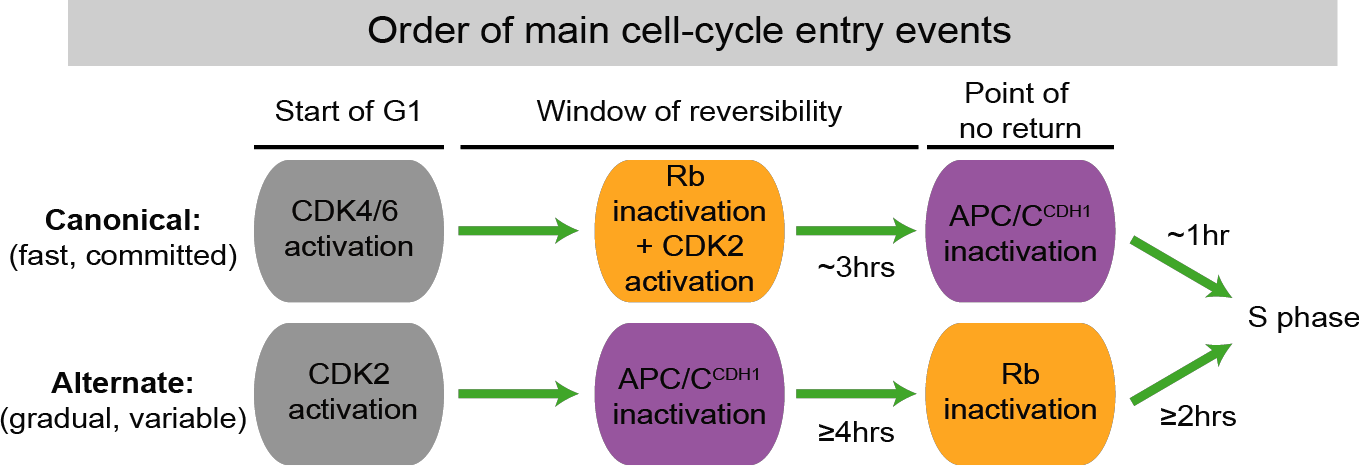
Working model.

Given the increasing number of usage for CDK4/6 inhibitors in cancer treatment and the potential of cyclin E-CDK2 mediated drug resistance^20–23^, our study also has therapeutic relevance. Our data showed that the alternate pathway is often marked by fluctuating cyclin E-CDK2 activity and is more sensitive to stress (Fig. 2b-d), and these unique features may offer opportunities for exploiting synthetic lethality in cancers. Finally, when considering that (1) >10 billion normal cells need to be replaced every day^50,51^ and (2) CDK4/6 inhibitors do not have toxicity issues common among other CDK inhibitors (the side effects are primarily hematopoietic)^52^, the use of the alternate pathway may explain why most normal tissues tolerate CDK4/6 inhibitors during cancer therapy.

## Acknowledgments

We thank M. Köberlin, L. Pack, and A. Cunningham for critical reading of the manuscript. We thank past and present members of the Meyer lab for reagents and discussions, Karlene Cimprich, James Ferrell, and Julien Sage for helpful discussions, and the Stanford Shared FACS facility for cell sorting. We also thank the laboratories of Richard Mort, Steven Cappell, Karlene Cimprich, and Thomas Wandless for providing the Fucci2a mice, constructs, cell lines, and chemicals. C.L. was supported by the NSF Graduate Research Fellowship, L.H.D. was supported by NIH training grant T32GM007365, and T.M. was supported by NIH grants GM118377, GM030179, and P50GM107615. Y.K., K.T., and M.M. were supported by JSPS 15H05949 “Resonance Bio” and 16H06280 “ABiS”. Further information and requests for resources and reagents should be directed to and will be fulfilled by the lead contact, Tobias Meyer.

## Conflict of Interests

The authors declare no competing interests.

## Materials and Methods

### General experimental setup

Cells were seeded to ensure 30% to 90% confluency throughout the experiment in 96 well plates. In the majority of the experiments, the same numbers of cells were plated for all conditions (one exception being the multigenerational experiment that involved 72 hours of live imaging, where the DMSO condition started off with 75% of the cells in the CDK4/6 inhibitor condition). In mitogen starvation experiments, cells were left in the starvation media for 48 hours after being washed at least once. Cells were then mitogen released with and without Palbociclib and refreshed every 24 hours unless indicated otherwise.

### Cell lines

All cell lines were acquired from ATCC unless noted otherwise. Cells were always grown in 37°C and 5% CO_2_ and between 30% and 80% confluency. Cells were provided fresh media at least once every three days. Dulbecco Modified Eagle medium (DMEM) and DMEM/F12 were acquired from ThermoFisher Scientific. MCF-10A (ATCC, #CRL-10317, human female) were cultured in phenol red-free DMEM/F12 supplemented with 5% horse serum, 20 ng/mL EGF, 10 μg/mL insulin, 500 μg/mL hydrocortisone, and 100 ng/mL cholera toxin. Starvation media consisted of the same growth media but instead of horse serum, insulin, and EGF, 0.3% BSA is added. Re-suspension media (used to inactivate trypsin) consisted of DMEM/F12 and 20% horse serum (protocol from the laboratory of Joan Brugge). All MCF-10A experiments were conducted on cells that went through <15 passages (passage 1 being receipt from ATCC), and cell line identity was confirmed through RNA-seq. BJ-5ta (ATCC, #CRL-4001, human male) were cultured in DMEM plus 10% FBS, 20% Medium 199 (ThermoFisher), and 0.02mg/ml hygromycin B. Starvation media consisted of DMEM, 0.1% BSA, and 0.02mg/ml hygromycin B. HS68 (ATCC, #CRL-1635, human male), passage 10, and WI-38 (ATCC, #CCL-75, human female), passage 5, were cultured in DMEM plus 10% FBS. MDA-MB-435 (ATCC, #HTB-129, human female) were cultured in Leibovitz’s L-15 medium (ATCC) plus 0.01mg/mL bovine insulin, 0.01mg/mL glutathione, and 10% FBS. Colo-205 (ATCC, #CCL-222, human male) was cultured in RPMI 1640 (ThermoFisher) with 10% FBS. U2OS (human female), acquired from the laboratory of Karlene Cimprich, and HeLa (ATCC, #CCL-2, human female) were cultured in DMEM plus 10% FBS. Panc 08.13 (ATCC, #CRL-2551, human male) were cultured in RPMI-1640 (ThermoFisher) plus 10 Units/mL human recombinant insulin, and 15% FBS. SCC-9 (ATCC, #CRL-1629, human male) were cultured in 1:1 mixture of DMEM and Ham’s F12 containing 1.2g/L sodium bicarbonate, 2.5mM L-glutamine, 15mM HEPES and 0.5mM sodium pyruvate plus 400ng/mL hydrocortisone and 10% FBS (protocol from ATCC).

### Stably transfected cell lines

All constructs were introduced into cells by lentiviral transduction. CSII-pEF-H2B-mTurquoise, CSII-pEF-DHB(aa994-1087)-mVenus, CSII-pEF-DHB(aa994-1087, serine to alanine mutant)-mCherry and CSII-pEF-mCherry-Geminin(aa1-110) were described previously^12,30,31^. Transduced cells were sorted on a Becton Dickinson Influx to obtain populations expressing the desired fluorescent reporters. DHFR-Myc^53^ was cloned into pCru5-IRES-puro and selected with 1 μg/ml puromycin (until the mock transfected cells all died).

### Chemical inhibitors, DNA damage, salt, H2O2, EdU treatment

The inhibitors used in this study were: CDK1 inhibitor RO3306 at 10μM (Sigma-Aldrich SML0569), CDK4/6 inhibitor PD0332991 (Palbociclib) at 1μM unless indicated otherwise (Selleck Chemicals S1116), CDK4/6 inhibitor LY2835219 (Abemaciclib) at 3μM (Selleck Chemicals S7158), CDK4/6 inhibitor LEE011 (Ribociclib) at 9μM (Selleck Chemicals S7440), and CDK1/2 inhibitor CAS 443798-55-8 at 3μM (EMD Biosciences #217714).

In experiments involving CDK4/6 inhibitor, cells were treated long term. In Rb phosphorylation experiments involving CDK1i and CDK1/2i, cells were treated for the last 20 minutes prior to fixation. In APC degron experiments involving CDK1/2i, cells were treated with CDK1/2i for 4 hours. In experiments involving neocarzinostatin (NCS) and aphidicolin, the concentrations and durations are indicated in the legends and figures. In experiments where hypertonic conditions are created, a 3M stock of salt in water (30x) is spiked in cells, the final media osmolality is ~440mOsm/kg. For oxidative stress, 200μM of H2O2 were applied. For assays involving EdU staining, cells were treated with 10μM EdU for ~15 minutes or 100μM for 5 minutes prior to fixation unless noted otherwise (cells in Figure 1c and Extended Data Figure 2c were treated with 100nM EdU for 24hrs).

### siRNA transfection

Cells were transfected using Dharmafect 1 (ThermoFisher) according to the manufacturer’s instructions. The following Dharmacon siRNAs were used: control siRNA (nontargeting #2), siGenome Human Myc (4609) set of four, Human siGenome Human CCNE1 (3213-09), siGenome Human CDKN1A (1026) set of four, siGenome FBXO5 (12434-01, 02, 03). siRNAs were used at a final concentration of 20nM. Cells were transfected 30 to 40 hours after starvation (the siRNAs were diluted in starvation media) and washed out when released with growth factors.

### c-Myc induction

Cells stably expressing DHFR-Myc were treated with 10μM Trimethoprim (TMP) at the time of release. TMP was refreshed every 24hrs along with the CDK4/6 inhibitor.

### Time-lapse microscopy

Images were taken on an IXMicro microscope (Molecular Devices). 10X objective (0.3 N.A.) used for live-cell imaging and fixed-cell immunofluorescence imaging. 20X objective (0.75 N.A.) with 2-by-2 pixel binning for imaging RNA FISH staining. For live-cell imaging, images were taken every 12 minutes, and total light exposure time was kept under 600ms for each time point. Cells were imaged in a humidified, 37°C chamber in 5% CO_2_.

### Tissue culture immunofluorescence

Tissue culture cells were fixed in 4% paraformaldehyde for 10 minutes, then blocked in PBS containing 10% FBS, 1% BSA, 0.1% TX-100, and 0.01% NaN3 for 1 hour at room temperature, and then stained overnight in 4ºC with antibodies for immunofluorescence (in blocking buffer). Alexa Fluor secondary goat antibodies (ThermoFisher) were applied 1:2000 for one hour at room temperature in blocking buffer, and then added with Hoechst 33342 (ThermoFisher) at 1:10000 for ten minutes at room temperature. Cells are stored in PBS when imaged. In the case where cells expressed fluorescent reporters that limited the use of fluorophores, cells were chemically bleached after fixation^54^. In the case of cyclin E1 staining, ice-cold methanol was applied for ten minutes instead of paraformaldehyde for fixation. In experiments where incorporated EdU signal is measured, the Click reaction is performed after blocking (if photobleaching is required, Click reaction also is performed after) following manufacturer’s protocol (Invitrogen, #C10269).

Antibodies used: Rb(p-S807/811) (Cell Signaling, #8516, 1:2500), Rb (Cell Signaling, #9309, 1:1000), Cyclin E (Santa Cruz, sc-247, 1:400), p21 (Cell Signaling, #2947, 1:250), geminin (Human Atlas, HPA049977, 1:500), c-Myc (Cell Signaling, #5605, 1:800), γH2A.X(S139) (Cell Signaling, #2577 1:800), 53BP1 (EMD Millipore MAB3802, 1:1000).

### RNA FISH

RNA in situ hybridization was carried out using the Affymetrix Quantigene ViewRNA ISH cell assay as specified in the user manual right after fixation (precedes Click reaction or immunofluorescence). Pre-made probes were designed to target E2F1 and EMI1. RNA in situ hybridization was carried out using the Affymetrix Quantigene ViewRNA ISH cell assay kit following manufacture’s protocol.

### Pre-extraction

Rb pre-extraction protocol was described previously^39^, but rather than trypsinizing and FACS, pre-extraction and fixation were performed with the 96-well plate on an ice block. Antibody used: Rb (BD Biosciences #554136, 1:250). When EdU needs to be incorporated, the incorporation precedes the permeabilization.

### Treatment, isolation, and immunofluorescence of mice small intestinal crypts

Fucci2a mice (expression of mCherry-hCdt1(30-120)-T2A-Venus-hGem(1-110), RDB13080, RIKEN)^40^ were made by crossing lox-stop-lox Fucci2a mice (Gt(ROSA)26Sor tm1(Fucci2aR)Jkn) with EIIa-Cre mice (B6.FVB-Tg (EIIa-cre) C5379Lmgd/J). ElIa-Cre mice were a gift from Mitinori Saitou, Kyoto University, Kyoto, Japan. The mice were dosed with 150mg/kg of Palbociclib (30mg/mL in sodium lactate buffer). Palbociclib was prepared fresh and administered orally via gavage. Mice were re-dosed after 24hrs and sacrificed after 48hrs. Immediately after sacrifice, the proximal, central, and distal regions of the small intestine were collected and fixed in 4% paraformaldehyde overnight in 4°C. Tissues were then washed and treated with 12% sucrose for 2hrs, 15% sucrose for 2hrs, and then 18% sucrose overnight (all sucrose steps conducted in 4°C). Tissues were then embedded in Optimal Cutting Temperature (O.C.T.) compound (Sakura Finetek Japan), and cryosectioned at 6μm thickness. Samples were then air-dried, washed, permeabilized/blocked (0.1% Triton X-100 and 1% BSA in TBST for 1hr in room temperature), and treated with primary antibody Rb(p-S807/811) (Cell Signaling, #8516, 1:1000) in blocking buffer at 4°C overnight. Alexa Fluor secondary goat anti-rabbit 647 (ThermoFisher) was applied at 1:2000 for one hour at room temperature in blocking buffer, and images were acquired immediately after using an FV1000/IX83 confocal microscope (Olympus). To avoid bleedthrough from the mCherry signal, phospho-Rb signal was collected at wavelengths of 690-790nm. Both female and male mice were used, and the mice age range from 6 weeks to 11 weeks. The animal protocols were reviewed and approved by the Animal Care and Use Committee of Kyoto University Graduate School of Medicine (No. 18086).

### Quantification and Statistical Analysis

Image analyses for live-cell and fix-cell assays have been described previously^12^. Derivation of APC/C^CDH1^ activity has been described previously^12^. In Extended Data Figure 5a, Figure 5, and Extended Data Figure 8, the time of APC/C^CDH1^ inactivation is moved earlier by 48 minutes to adjust for the mCherry maturation time^12^. Analysis for RNA FISH has been described previously^29^. In this manuscript, the FISH, γH2A.X(S139), and 53BP1 puncta counts were quantified by counting each pixel above a threshold (same threshold is used within an experiment). Information about replicates and error bars can be found in the figure legends. In box plots, the central line indicates median, the edges of the box denote the quartiles, and the whiskers extend to the farthest points that are not outliers (defined as 1.5 times the interquartile range). Automated-pipeline scripts performed all analyses. In analyses where only certain populations of cells are analyzed (for example, G0/G1 cells), the gating criteria are described in the figures, figure legends, or results section. In experiments where significance was derived, unpaired two-sample t-tests were performed. In Extended Data Figure 8a, the time denoted by the gray box is estimated.

Experiments with MCF-10A and BJ-5ta are excluded when cells are of suboptimal confluency and/or unhealthy (<30% of control cells in S/G2 24hrs after mitogen release or when asynchronously cycling).

Crypts were selected for analysis based on the following criteria: (i) contains at least ten cells expressing the sensors, (ii) shaped like the letter U, and (iii) contains post-mitotic cells as negative controls in the bottom of the crypt. Cells were segmented manually (by merging the three nuclear channels: geminin-fragment, Cdt1-fragment, and phospho-Rb) and median nuclear intensities were then calculated. To calculate the thresholds for determining APC/C-degron positive and phospho-Rb positive cells, the signals of post-mitotic cells from the same crypt were averaged and multiplied by a small factor. The same unbiased approach for threshold determination is applied to every image.

## Data and Code Availability

All code and data are available from the corresponding author upon reasonable request.

## Author Contributions

Conceptualization, C.L. and T.M.; Methodology, C.L., and M.C.; Software: C.L. and M.C.; Formal Analysis: C.L.; Investigation, C.L., Y.K., M.C., L.H.D., and Y.F.; Resources: C.L., H.W.Y., K.T., M.M., and T.M.; Writing, C.L., and T.M.; Visualization: C.L.; Supervision: T.M.; Funding Acquisition: C.L., K.T., M.M., T.M.

## References and Notes

1. Matson, J. P. & Cook, J. G. Cell cycle proliferation decisions: the impact of single cell analyses. FEBS J. 284, 362–375 (2017).

2. Fisher, R. P. Getting to S: CDK functions and targets on the path to cell-cycle commitment. F1000Research 5, 2374 (2016).

3. Sherr, C. J. G1 phase progression: Cycling on cue. Cell 79, 551–555 (1994).

4. Weinberg, R. A. The retinoblastoma protein and cell cycle control. Cell 81, 323–330 (1995).

5. Nevins, J. R. The Rb/E2F pathway and cancer. Hum. Mol. Genet. 10, 699–703 (2001).

6. Bracken, A. P., Ciro, M., Cocito, A. & Helin, K. E2F target genes: unraveling the biology. Trends Biochem. Sci. 29, 409–417 (2004).

7. Hsu, J. Y., Reimann, J. D. R., Sørensen, C. S., Lukas, J. & Jackson, P. K. E2F-dependent accumulation of hEmi1 regulates S phase entry by inhibiting APCCdh1. Nat. Cell Biol. 4, 358–366 (2002).

8. Salazar-Roa, M. & Malumbres, M. Fueling the Cell Division Cycle. Trends Cell Biol. 27, 69–81 (2017).

9. Peters, J.-M. The anaphase promoting complex/cyclosome: a machine designed to destroy. Nat. Rev. Mol. Cell Biol. 7, 644–656 (2006).

10. Yamada, M. et al. ATR-Chk1-APC/CCdh1-dependent stabilization of Cdc7-ASK (Dbf4) kinase is required for DNA lesion bypass under replication stress. Genes Dev. 27, 2459–72 (2013).

11. Barr, A. R., Heldt, F. S., Zhang, T., Bakal, C. & Novák, B. A Dynamical Framework for the All-or-None G1/S Transition. Cell Syst. 2, 27–37 (2016).

12. Cappell, S. D., Chung, M., Jaimovich, A., Spencer, S. L. & Meyer, T. Irreversible APCCdh1 Inactivation Underlies the Point of No Return for Cell-Cycle Entry. Cell 166, 167–180 (2016).

13. Cappell, S. D. et al. EMI1 switches from being a substrate to an inhibitor of APC/C^CDH1^ to start the cell cycle. Nature 558, 313–317 (2018).

14. Kozar, K. et al. Mouse Development and Cell Proliferation in the Absence of D-Cyclins. Cell 118, 477–491 (2004).

15. Liu, L. et al. G1 cyclins link proliferation, pluripotency and differentiation of embryonic stem cells. Nat. Cell Biol. 19, 177–188 (2017).

16. Alevizopoulos, K., Vlach, J., Hennecke, S. & Amati, B. Cyclin E and c-Myc promote cell proliferation in the presence of p16INK4a and of hypophosphorylated retinoblastoma family proteins. EMBO J. 16, 5322–33 (1997).

17. Prall, O. W., Rogan, E. M., Musgrove, E. A., Watts, C. K. & Sutherland, R. L. c-Myc or cyclin D1 mimics estrogen effects on cyclin E-Cdk2 activation and cell cycle reentry. Mol. Cell. Biol. 18, 4499–508 (1998).

18. Lukas, J. et al. Cyclin E-induced S phase without activation of the pRb/E2F pathway. Genes Dev. 11, 1479–92 (1997).

19. Jiang, H., Chou, H. S. & Zhu, L. Requirement of cyclin E-Cdk2 inhibition in p16(INK4a)-mediated growth suppression. Mol. Cell. Biol. 18, 5284–90 (1998).

20. Taylor-Harding, B. et al. Cyclin E1 and RTK/RAS signaling drive CDK inhibitor resistance via activation of E2F and ETS. Oncotarget 6, 696–714 (2015).

21. Herrera-Abreu, M. T. et al. Early Adaptation and Acquired Resistance to CDK4/6 Inhibition in Estrogen Receptor-Positive Breast Cancer. Cancer Res. 76, 2301–13 (2016).

22. Dean, J. L., Thangavel, C., McClendon, A. K., Reed, C. A. & Knudsen, E. S. Therapeutic CDK4/6 inhibition in breast cancer: key mechanisms of response and failure. Oncogene 29, 4018–4032 (2010).

23. Patel, P., Tsiperson, V., Gottesman, S. R. S., Somma, J. & Blain, S. W. Dual Inhibition of CDK4 and CDK2 via Targeting p27 Tyrosine Phosphorylation Induces a Potent and Durable Response in Breast Cancer Cells. Mol. Cancer Res. 16, 361–377 (2018).

24. Sakaue-Sawano, A. et al. Visualizing Spatiotemporal Dynamics of Multicellular Cell-Cycle Progression. Cell 132, 487–498 (2008).

25. Fry, D. W. et al. Specific inhibition of cyclin-dependent kinase 4/6 by PD 0332991 and associated antitumor activity in human tumor xenografts. Mol Cancer Ther 3, (2004).

26. Jorda, R. et al. How Selective Are Pharmacological Inhibitors of Cell-Cycle-Regulating Cyclin-Dependent Kinases? J. Med. Chem. 61, 9105–9120 (2018).

27. Barr, A. R. et al. DNA damage during S-phase mediates the proliferation-quiescence decision in the subsequent G1 via p21 expression. Nat. Commun. 8, 14728 (2017).

28. Arora, M., Moser, J., Phadke, H., Basha, A. A. & Spencer, S. L. Endogenous Replication Stress in Mother Cells Leads to Quiescence of Daughter Cells. Cell Rep. 19, 1351–1364 (2017).

29. Yang, H. W., Chung, M., Kudo, T. & Meyer, T. Competing memories of mitogen and p53 signalling control cell-cycle entry. Nature 549, 404–408 (2017).

30. Hahn, A. T., Jones, J. T. & Meyer, T. Quantitative analysis of cell cycle phase durations and PC12 differentiation using fluorescent biosensors. Cell Cycle 8, 1044–1052 (2009).

31. Spencer, S. L. et al. The Proliferation-Quiescence Decision Is Controlled by a Bifurcation in CDK2 Activity at Mitotic Exit. Cell 155, 369–383 (2013).

32. Schwarz, C. et al. A Precise Cdk Activity Threshold Determines Passage through the Restriction Point. Mol. Cell 69, 253–264.e5 (2018).

33. Merrick, K. A. et al. Distinct Activation Pathways Confer Cyclin-Binding Specificity on Cdk1 and Cdk2 in Human Cells. Mol. Cell 32, 662–672 (2008).

34. Daigh, L. H., Liu, C., Chung, M., Cimprich, K. A. & Meyer, T. Stochastic Endogenous Replication Stress Causes ATR-Triggered Fluctuations in CDK2 Activity that Dynamically Adjust Global DNA Synthesis Rates. Cell Syst. (2018). doi:10.1016/j.cels.2018.05.011

35. El-Deiry, W. S. et al. WAF1, a potential mediator of p53 tumor suppression. Cell 75, 817–825 (1993).

36. Hittelman, W. N. & Pollard, M. Induction and repair of DNA and chromosome damage by neocarzinostatin in quiescent normal human fibroblasts. Cancer Res. 42, 4584–90 (1982).

37. Narasimha, A. M. et al. Cyclin D activates the Rb tumor suppressor by mono-phosphorylation. Elife 3, (2014).

38. Zhang, H. S. & Dean, D. C. Rb-mediated chromatin structure regulation and transcriptional repression. Oncogene 20, 3134–3138 (2001).

39. Håland, T. W., Boye, E., Stokke, T., Grallert, B. & Syljuåsen, R. G. Simultaneous measurement of passage through the restriction point and MCM loading in single cells. Nucleic Acids Res. 43, e150–e150 (2015).

40. Mort, R. L. et al. *Fucci2a*: A bicistronic cell cycle reporter that allows Cre mediated tissue specific expression in mice. Cell Cycle 13, 2681–2696 (2014).

41. Gehart, H. & Clevers, H. Tales from the crypt: new insights into intestinal stem cells. Nat. Rev. Gastroenterol. Hepatol. 16, 19–34 (2019).

42. Ferrell, J. E. Tripping the switch fantastic: how a protein kinase cascade can convert graded inputs into switch-like outputs. Trends Biochem. Sci. 21, 460–466 (1996).

43. Ferrell, J. E. Self-perpetuating states in signal transduction: positive feedback, double-negative feedback and bistability. Curr. Opin. Cell Biol. 14, 140–148 (2002).

44. Tyson, J. J., Chen, K. C. & Novak, B. Sniffers, buzzers, toggles and blinkers: dynamics of regulatory and signaling pathways in the cell. Curr. Opin. Cell Biol. 15, 221–231 (2003).

45. Di Fiore, B. & Pines, J. Emi1 is needed to couple DNA replication with mitosis but does not regulate activation of the mitotic APC/C. J. Cell Biol. 177, 425–37 (2007).

46. Machida, Y. J. & Dutta, A. The APC/C inhibitor, Emi1, is essential for prevention of rereplication. Genes Dev. 21, 184–94 (2007).

47. Malumbres, M. et al. Mammalian Cells Cycle without the D-Type Cyclin-Dependent Kinases Cdk4 and Cdk6. Cell 118, 493–504 (2004).

48. Sage, J., Miller, A. L., Pérez-Mancera, P. A., Wysocki, J. M. & Jacks, T. Acute mutation of retinoblastoma gene function is sufficient for cell cycle re-entry. Nature 424, 223–228 (2003).

49. Rossi, A. et al. Genetic compensation induced by deleterious mutations but not gene knockdowns. Nature 524, 230–233 (2015).

50. Nagata, S. Apoptosis and Clearance of Apoptotic Cells. Annu. Rev. Immunol. 36, 489–517 (2018).

51. Elliott, M. R. & Ravichandran, K. S. The Dynamics of Apoptotic Cell Clearance. Dev. Cell 38, 147–160 (2016).

52. Sánchez-Martínez, C., Gelbert, L. M., Lallena, M. J. & de Dios, A. Cyclin dependent kinase (CDK) inhibitors as anticancer drugs. Bioorg. Med. Chem. Lett. 25, 3420–3435 (2015).

53. Iwamoto, M., Björklund, T., Lundberg, C., Kirik, D. & Wandless, T. J. A General Chemical Method to Regulate Protein Stability in the Mammalian Central Nervous System. Chem. Biol. 17, 981–988 (2010).

54. Lin, J.-R., Fallahi-Sichani, M. & Sorger, P. K. Highly multiplexed imaging of single cells using a high-throughput cyclic immunofluorescence method. Nat. Commun. 6, 8390 (2015).

