## Supplemental Figures for "A parallel cell-cycle entry pathway with inverted G1 signaling and alternate point of no return"

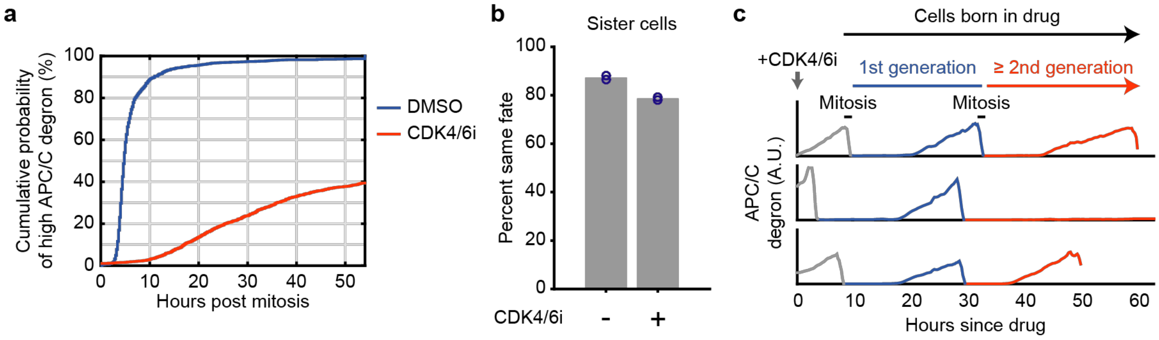


**Extended Data Figure 1. Related to Figure 1**

**a,** CDF of asynchronously cycling cells inactivating APC/C$^{CDH1}$ after mitosis. 1 of n=2 biological replicates. **b,** Percent sister cells with the same fate as assayed by APC/C inactivation. Each dot represents a biological replicate. **c,** Example live-cell traces demonstrating separation of first and subsequent generations in MCF-10A cells. Example cells were born into CDK4/6 inhibitor and drug was refreshed every 24hrs.

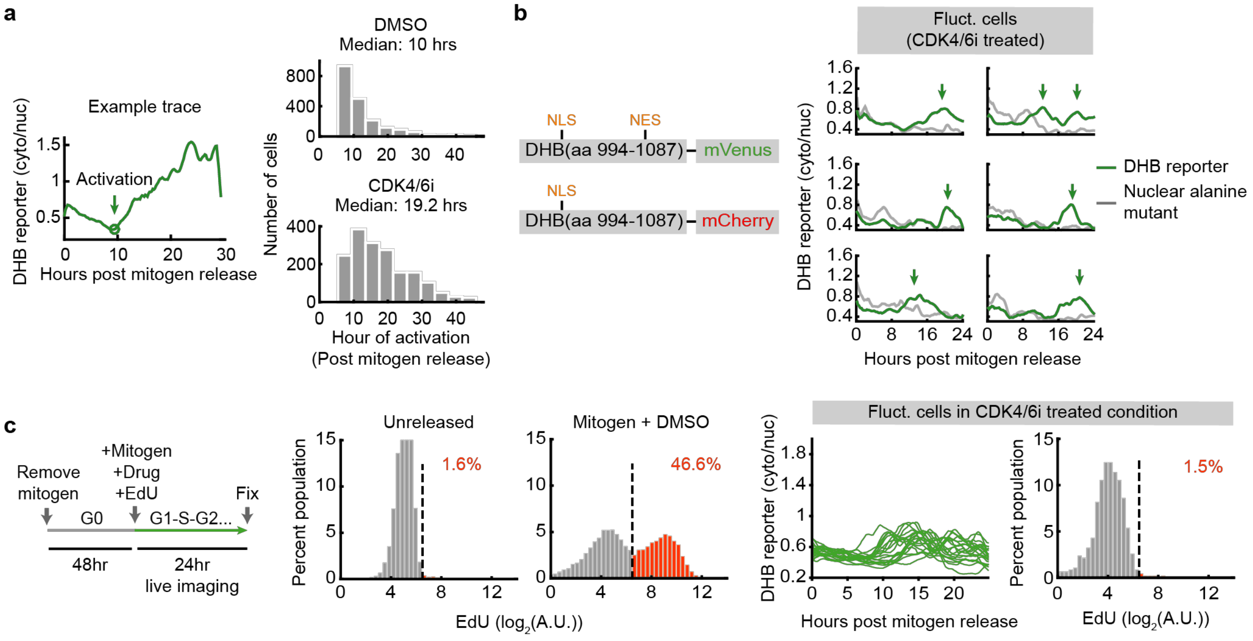


**Extended Data Figure 2. Related to Figure 2. Delayed and fluctuating cyclin E/A-CDK activity in CDK4/6 inhibitor-treated cells**

**a,** Left: Example trace of cyclin E/A-CDK activity with circle denoting point of activation. Right: Distributions of activation time in cells coming out of mitogen release. **b,** Left: Co-transfected constructs. Right: Example traces of cells expressing both DHB reporter (green) and nuclear DHB-alanine-mutant reporter (gray). Green arrows denote fluctuations not due to analysis noise. **c,** Left: cells were mitogen-released with EdU to assay for S-phase entry. Middle two histograms: negative and positive controls for determining EdU threshold. Right: EdU incorporation in cyclin E/A-CDK fluctuating cells (<0.7 at last time point, exceeded 0.7 at some point between 8 and 24hrs after mitogen release). 20 fluctuating cells plotted for the traces, 474 fluctuating cells total.

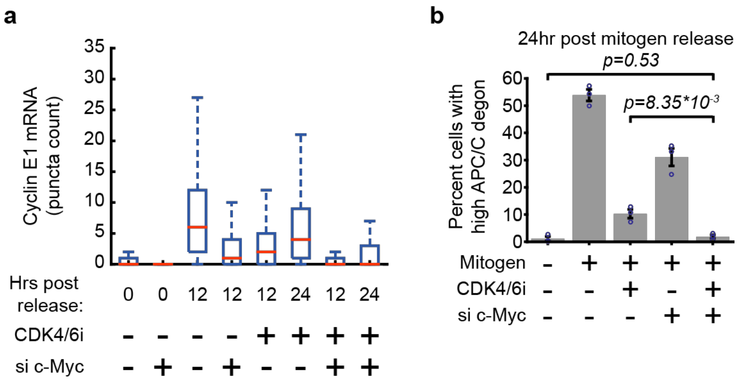


**Extended Data Figure 3. Related to Figure 2. c-Myc is necessary for APC/C$^{CDH1}$ inactivation in CDK4/6 inhibitor-treated cells**

**a,** Cyclin E1 mRNA puncta count in cells mitogen released with and without CDK4/6 inhibitor and si c-Myc (from left to right, n=5992, 8133, 10381, 9584, 8628, 6132, 9535, and 4681 cells). **b,** Cells with and without CDK4/6 inhibitor and si c-Myc were fixed 24hrs after mitogen release. Error bar: SEM from 3 biological replicates; p-values calculated using two-sided, two-sample t-tests.

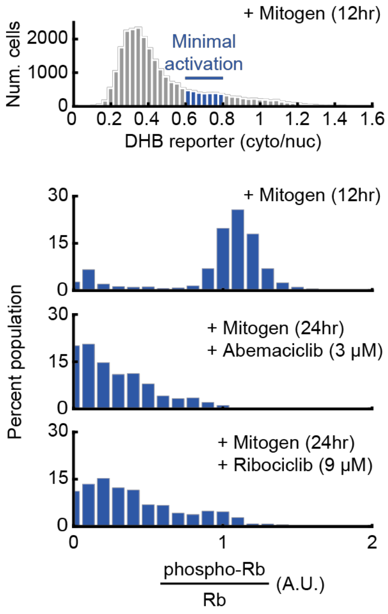


**Extended Data Figure 4. Related to Figure 3. Cyclin E/A-CDK activation without phosphorylated Rb in cells treated with different CDK4/6 inhibitors**

Cells expressing the cyclin E/A-CDK activity reporter and APC/C degron reporter were mitogen-released, fixed after 12hrs (DMSO) and 24hrs (Abemaciclib at 3µM and Ribociclib at 9µM), and stained for Rb and phospho-Rb(S807/S811). G1 cells with recently activated cyclin E/A-CDK activity were analyzed. n= 25622, 29553, and 28693 cells for DMSO, Abemaciclib, and Ribociclib conditions, respectively.

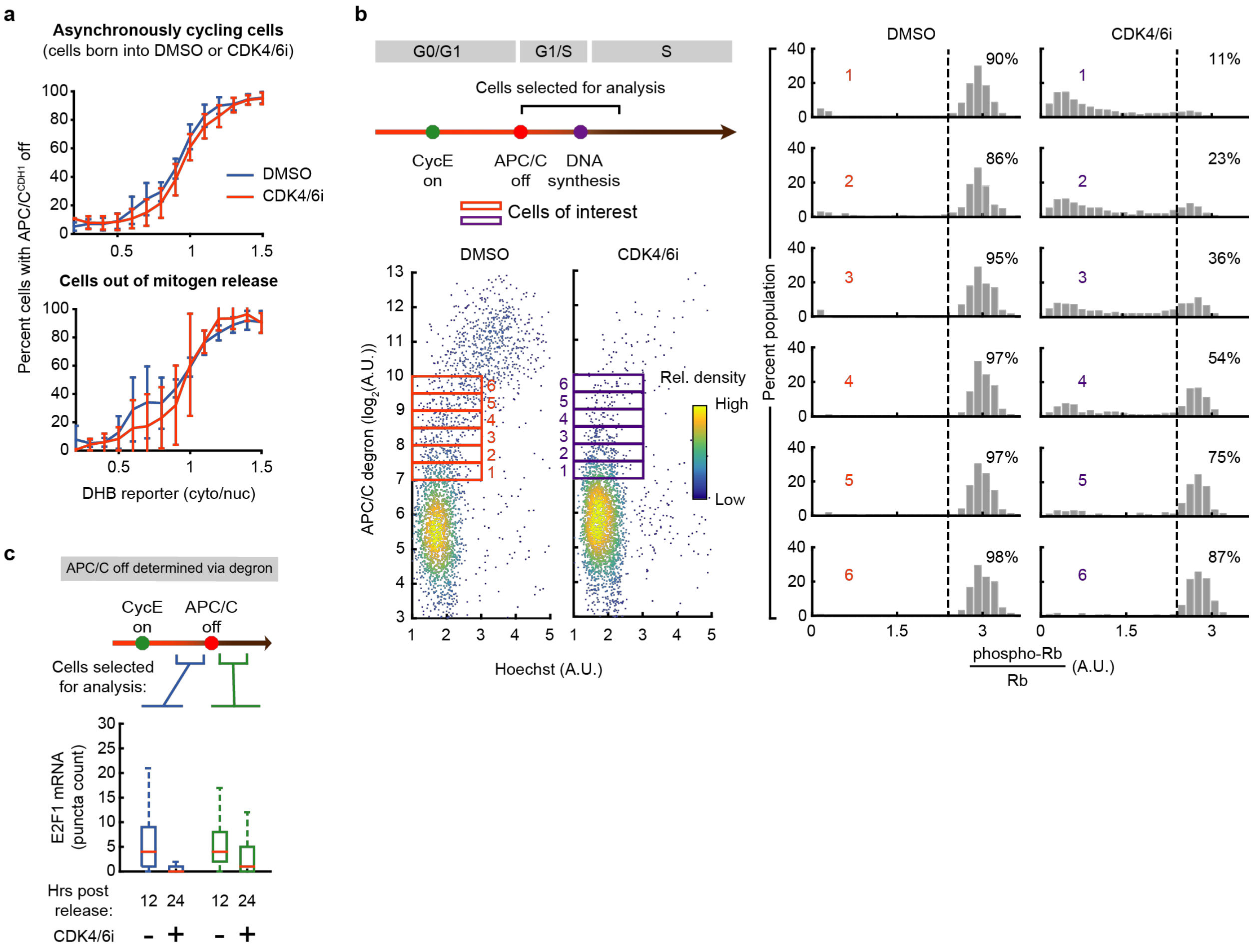


**Extended Data Figure 5. Related to Figure 4. Rb phosphorylation and E2F activation occurs after APC/C$^{CDH1}$ inactivation in CDK4/6-inhibited cells**

**a,** The percentages of cells with APC/C$^{CDH1}$ inactivated were determined for each bin of the cyclin E/A-CDK activity. Top: Cells born into DMSO or CDK4/6 inhibitor. Error bars denote standard deviation of 3 biological replicates. Bottom: Cells mitogen-released with DMSO or CDK4/6 inhibitor. Error bars denote standard deviation of 4 biological replicates. **b,** Asynchronously cycling cells born into DMSO or 1μM CDK4/6 inhibitor and have recently inactivated APC/C$^{CDH1}$ were analyzed for phospho-Rb(S807/S811) signal (3000 cells plotted for scatter plots. >180 cells per histogram for DMSO, >350 cells per histogram for CDK4/6 inhibitor; 1 of n=2 biological replicates). **c,** E2F1 mRNA in cells released with or without CDK4/6 inhibitor. High cyclin E/A-CDK activity defined as ratio of signal>0.7. APC/C$^{CDH1}$ inactivation determined via the degron-based activity reporter. From left to right, n=8747, 4063, 1961, and 189 cells). 1 of n=2 biological replicates.

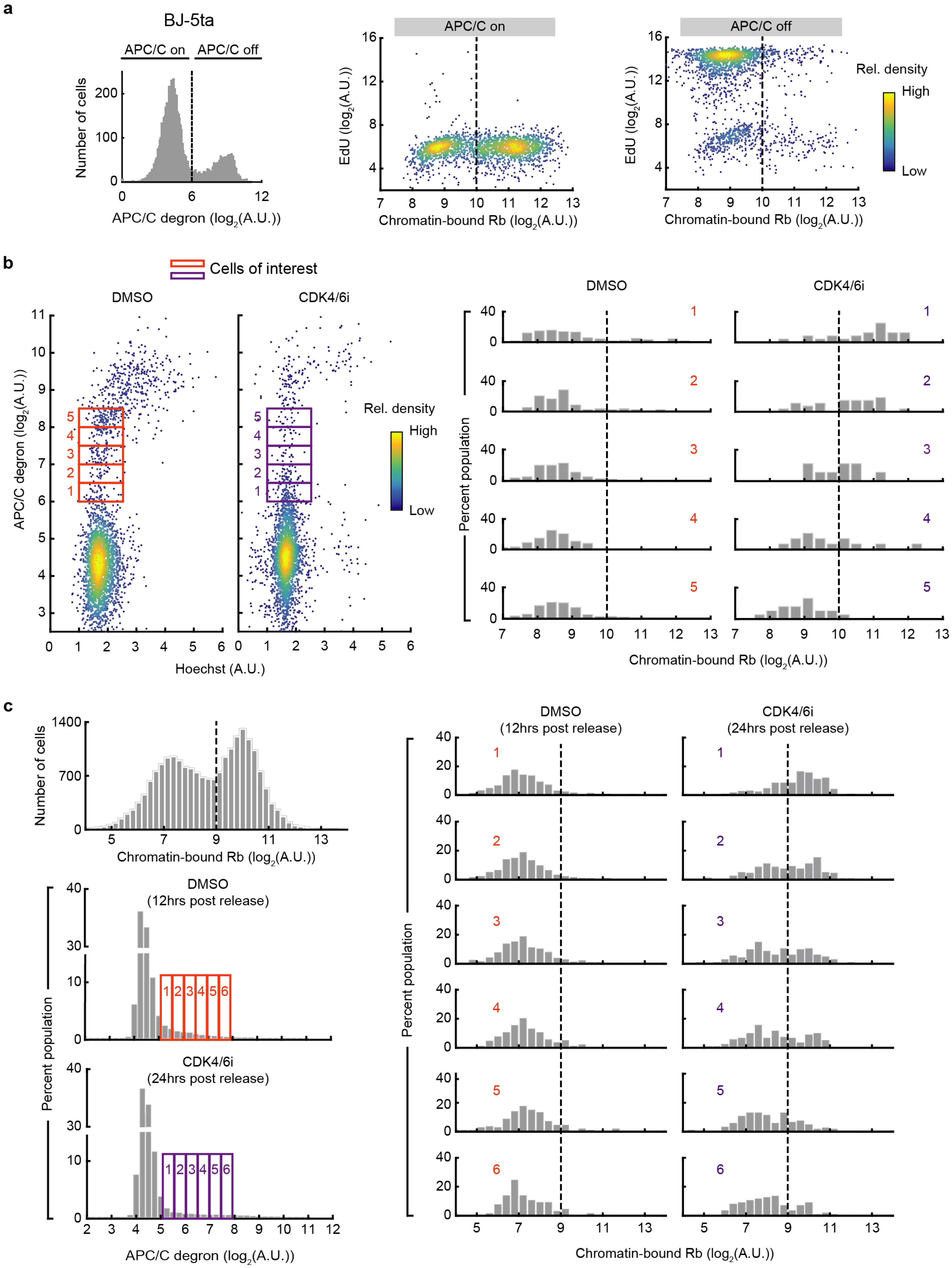
a
BJ-5ta
APC/C on
APC/C off
Number of cells
APC/C degron ($log_2$(A.U.))
EdU ($log_2$(A.U.))
Chromatin-bound Rb ($log_2$(A.U.))
Rel. density
High
Low
b
Cells of interest
DMSO
CDK4/6i
APC/C degron ($log_2$(A.U.))
Hoechst (A.U.)
Percent population
Chromatin-bound Rb ($log_2$(A.U.))
c
Number of cells
Chromatin-bound Rb ($log_2$(A.U.))
DMSO
(12hrs post release)
CDK4/6i
(24hrs post release)
Percent population
APC/C degron ($log_2$(A.U.))

**Extended Data Figure 6. Related to Figure 4. Validation and quantification of chromatin-bound Rb in CDK4/6-inhibited cells.**

**a,** BJ-5ta cells were treated with CDK4/6 inhibitor for 96hrs and 100μM EdU for the final 5min, then permeabilized/pre-extracted, fixed, and separated into APC/C on and APC/C off. Cells were then assayed for EdU and Rb signals (2000 cells plotted). Drugs were refreshed every 24hrs. **b,** Cells from **a** that recently inactivated $APC/C^{CDH1}$ were analyzed for pre-extracted Rb signal. **c,** MCF-10A cells were mitogen-released for 12hrs (with DMSO) or 24hrs (with CDK4/6 inhibitor), pre-extracted, and fixed. Cells that have recently inactivated $APC/C^{CDH1}$ were analyzed for pre-extracted Rb signal. 1 of n=2 biological replicates.

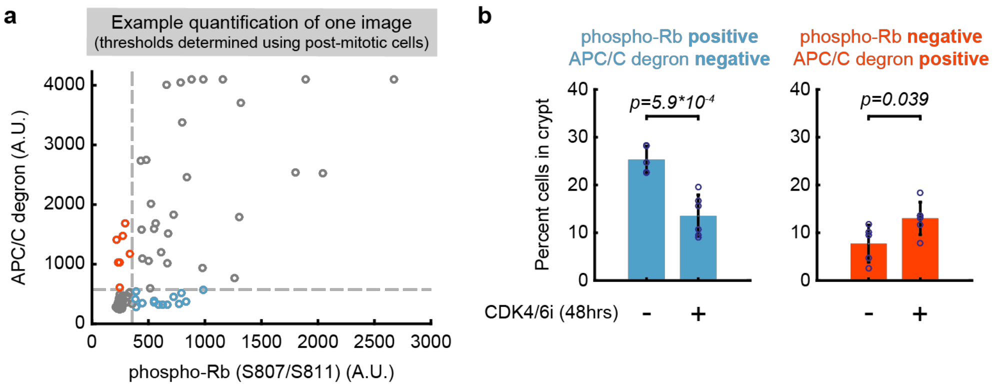


**Extended Data Figure 7. Related to Figure 4. Quantitation of small intestinal crypt cells**

**a,** Example scatter plot demonstrating the quantification process. Thresholds determined unbiasedly using post-mitotic cells, which have low phospho-Rb and APC/C degron signals. **b,** Quantification of percent cells with high phospho-Rb, low APC/C degron signal and vice versa. n=5 mice for control and n=6 mice for CDK4/6 inhibitor. Error bars: standard deviation. At least 100 intestinal crypt cells are quantified per mouse. p-values calculated using two-sided, two-sample t-tests.

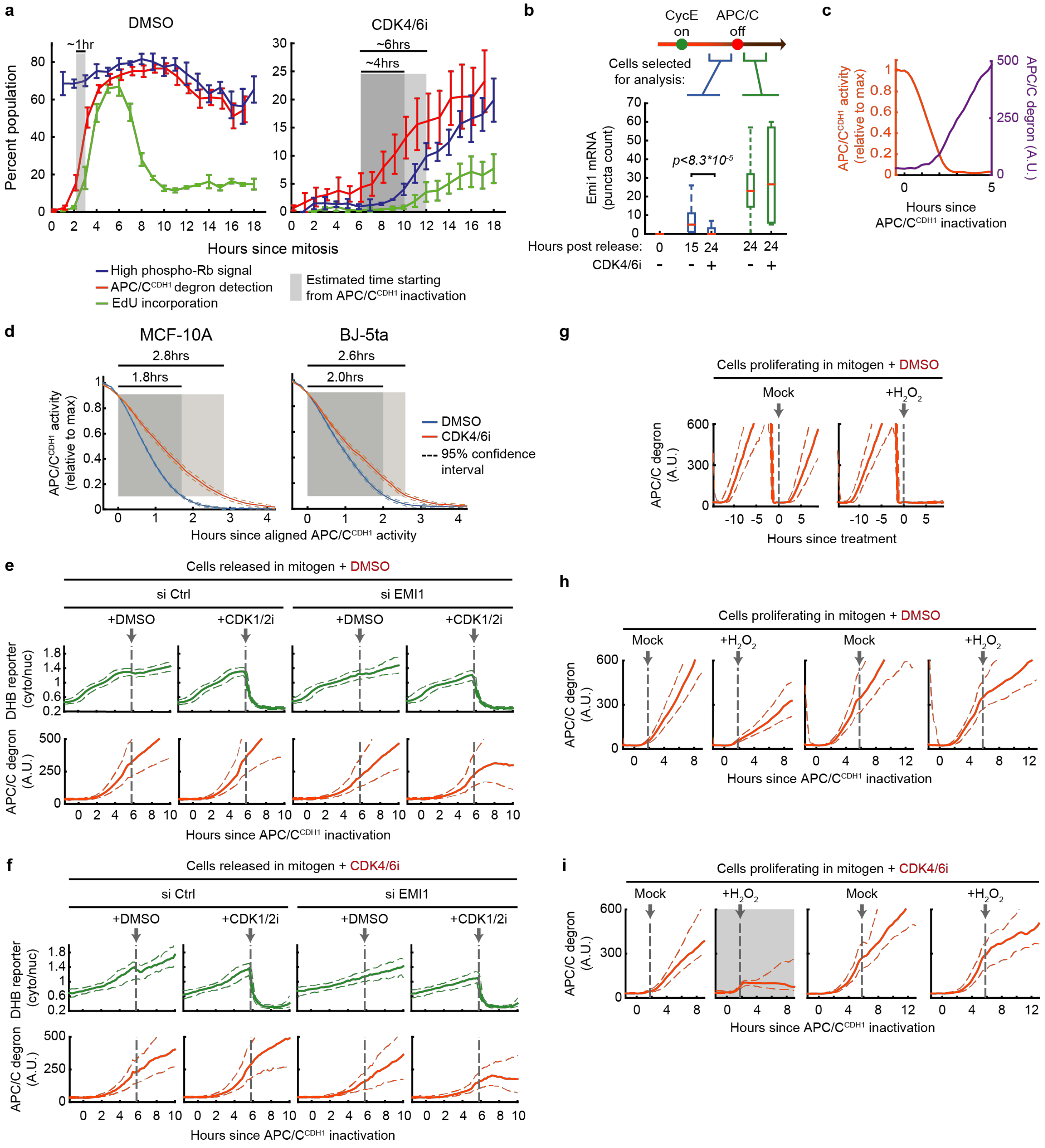


**Extended Data Figure 8. Related to Figure 5. Emi1 confers irreversibility to APC/C$^{CDH1}$ inactivation in the canonical and alternate pathway**

**a,** Cells born into DMSO or CDK4/6 inhibitor were aligned at the time of birth. For each hour after mitosis, the percent cells with high phospho-Rb(S807/S811) signal, high APC/C degron, and high EdU signal (after 15min of 10μM EdU treatment) were calculated. Error bars: SEM from 3 biological replicates. **b,** EMI1 mRNA in cells that were mitogen-released with or without 1μM CDK4/6 inhibitor. High cyclin E/A activity defined as reporter ratio>0.7. p-values calculated using two-sided, two-sample t-tests (n=44 and 95 cells for DMSO and CDK4/6 inhibitor conditions, respectively); 1 of n=3 biological replicates. **c,** Example of APC/C degron conversion to APC/$C^{CDH1}$ activity. **d,** Calculated APC/$C^{CDH1}$ inactivation kinetics in cells born into DMSO or CDK4/6 inhibitor. Shaded boxes denote 90% to 10% activity. Shown are the means with 95% confidence intervals (MCF-10A: n=3357 cells in DMSO and 601 cells in CDK4/6 inhibitor; BJ-5ta: n=1258 cells in DMSO and 322 cells in CDK4/6 inhibitor). **e-f,** Cells that were mitogen released into DMSO or CDK4/6 inhibitor and have inactivated APC/$C^{CDH1}$ were treated with 3μM CDK1/2i. Cells pooled from 4 biological replicates (Dashed lines denote $25^{th}$ and $75^{th}$ percentile. >100 cells per condition). **g-i,** Cells born into DMSO or CDK4/6 inhibitor and have inactivated APC/$C^{CDH1}$ (with the exception of **g**, where it is just cells born into DMSO or CDK4/6 inhibitor) were treated with 200μM $H_2O_2$. Dashed lines denote $25^{th}$ and $75^{th}$ percentile. >25 cells per condition. In **h-i**, to enrich for Rb inactivated cells in the 6hrs post APC/$C^{CDH1}$ inactivation conditions, only cells with cyclin E/A-CDK>0.8 right before treatment were analyzed (threshold determined from Figure 3e). Gray box denotes conditions where cells re-activated APC/$C^{CDH1}$.

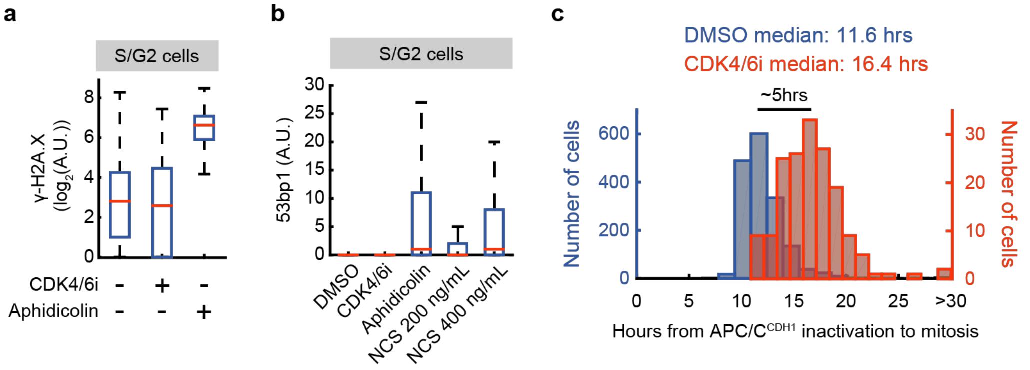


**Extended Data Figure 9. Related to Figure 5. Cells proliferating without CDK4/6 activity have a viable S phase**

**a,** Box plots of phospho-γH2A.X(S139) puncta count in S/G2 cells. Cells were treated with 1µM CDK4/6 inhibitor or 1µM aphidicolin (positive control) for 48hrs, drugs were refreshed at 24hrs (10421, 2319, 3850 cells for DMSO, CDK4/6 inhibitor, and aphidicolin, respectively). **b,** 53BP1 puncta count in S/G2 cells. Cells were treated with 1µM CDK4/6 inhibitor for 48hrs. Positive controls: 1µM aphidicolin (48hrs) and NCS (15min pulse 24hrs before fixation). Drugs were refreshed at 24hrs and only S/G2 cells were quantified (10223, 3406, 1931, 9326, 6619 cells for DMSO, CDK4/6 inhibitor, aphidicolin, NCS low, and NCS high, respectively). **c,** Time from APC/C$^{CDH1}$ inactivation to anaphase in cells that were born into DMSO or CDK4/6 inhibitor.
